# Elys deficiency constrains Kras-driven tumour burden by amplifying oncogenic stress

**DOI:** 10.1101/2021.08.25.457580

**Authors:** Kimberly J Morgan, Karen Doggett, Fan-Suo Geng, Lachlan Whitehead, Kelly A Smith, Benjamin M Hogan, Cas Simons, Gregory J Baillie, Ramyar Molania, Anthony T Papenfuss, Thomas E Hall, Elke A Ober, Didier Y R Stainier, Zhiyuan Gong, Joan K Heath

## Abstract

The nucleoporin ELYS, encoded by *AHCTF1*, is a large multifunctional protein with essential roles in nuclear pore assembly and mitosis. Using a zebrafish model of hepatocellular carcinoma, in which the expression of an inducible mutant *kras* transgene *(kras^G12V^)* drives hepatocyte-specific hyperplasia and liver enlargement, we show that reducing *ahctf1* gene dosage by 50% markedly shrinks tumour burden, while non-hyperplastic tissues are unaffected. We demonstrate that *ahctf1* heterozygosity impairs nuclear pore formation, mitotic spindle assembly and chromosome segregation, leading to DNA damage and activation of TP53-dependent and independent mechanisms of cell death and cell cycle arrest. This selective vulnerability of cancer cells to mild disruption of Elys function uncovers a novel synthetic lethal interaction between *ahctf1* and *kras* mutations that could be exploited therapeutically. Heterozygous expression of both *ahctf1* and *ranbp2*, or treatment of heterozygous *ahctf1* larvae with the nucleocytoplasmic transport inhibitor, Selinexor, completely blocked *kras^G12V^*-driven hepatocyte hyperplasia, revealing promising avenues for combinatorial treatments.

## INTRODUCTION

Synthetic lethality is the term used to describe the death of cells in response to co-existing disruptions in two different genes, neither of which is lethal alone. The phenomenon has emerged as a promising tool for cancer drug development^1^. The advantage of the approach lies in its capacity to induce the death of a vulnerable cell population, such as oncogene-expressing cancer cells, while healthy cells are unaffected. The approach has been validated in the clinic by the use of poly(adenosine diphosphate [ADP]-ribose) polymerase (PARP) inhibitors to treat tumours carrying mutations in the breast cancer susceptibility genes, *BRCA1*/*BRAC2*^2^, and its success has fuelled the search for other clinically relevant pairwise combinations. Particular effort has been directed towards identifying genes whose partial loss of function confers synthetic lethality in cancer cells containing hitherto ‘undruggable’ oncogene targets, such as mutant *KRAS*^3,4^. In this paradigm, the interacting gene is not mutated in cancer nor oncogenic in its own right; however, its function is essential to maintain the tumourigenic state, inspiring the concept of non-oncogene addiction^5^.

In this paper, we tested whether *AHCTF1* exhibits the properties of a mutant *KRAS* synthetic lethal interacting gene. *AHCTF1* encodes ELYS, a 252 kDa multidomain nucleoporin that was first discovered in mouse development where it was shown to be required for the proliferation and survival of inner mass cells^6^. Since then, ELYS has been studied in many model systems, including in a zebrafish development mutant (ti262)^7^, where we and others showed that homozygous inheritance of an ENU-induced nonsense mutation in *ahctf1* disrupted nuclear pore formation and caused catastrophic levels of cell death in highly proliferative cell compartments, such as the intestinal epithelium, while relatively quiescent tissues survived and remained healthy^8,9^.

As well as being a structural component of nuclear pore complexes (NPCs), ELYS is essential for their formation at the end of mitosis^10–12^. NPCs are huge (110 MDa) multi-subunit complexes comprising at least 30 different nucleoporins (Nups) in an octameric array^13^. They form cylindrical channels in the nuclear envelope (NE) and regulate nucleocytoplasmic transport and intracellular localisation of large (>40 kDa) molecules. In the absence of ELYS, the NE is re-built at the end of mitosis, but this occurs without NPC formation, precluding mRNAs and many proteins from moving in and out of the nucleus. By providing a platform for NPC re-assembly at the end of mitosis (Fig. 1a,b), ELYS restores nucleocytoplasmic trafficking, and thereby facilitates entry into the nucleus of proteins such as G1 and S phase cyclin-dependent kinases that are required for DNA replication and cell cycle progression (Fig. 1c). Also in G1/S phase, ELYS regulates transcription and DNA replication by reversible binding to components of the SWI/SNF 80 chromatin remodeller complex, PBAP, and the DNA replication origin licensing system, MCM2-7, respectively (Fig. 1d-f)^11,12,14,15^.

**Fig. 1.**
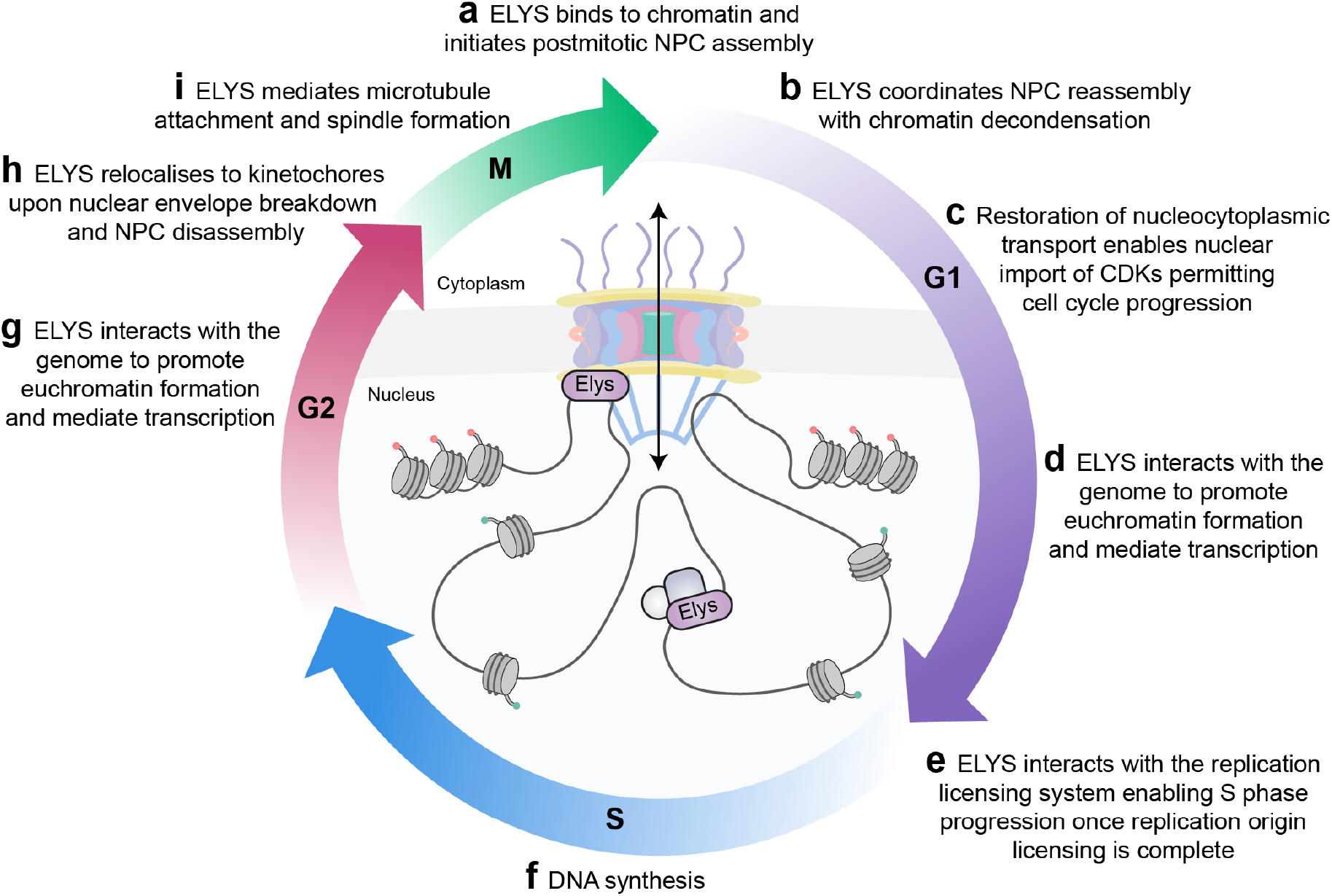
Elys plays critical roles at several stages of the cell cycle. Elys is a central player in multiple diverse cellular processes far beyond its canonical role in nuclear pore re-assembly at the end of mitosis **(a, b)**. **c.** Elys restores nucleocytoplasmic trafficking allowing import of cyclin-dependent kinases (CDKs) required for cell cycle progression. **d-g.** Elys plays an active role in genome regulation through its direct interactions with components of large molecular machineries including chromatin remodellers such as PBAP that favour transcription and the MCM2-7 components of the DNA replication origin licensing system. **h.** After cell growth in G2, the nuclear envelope breaks down and NPCs are disassembled upon entry into mitosis. **i.** Elys relocalizes to kinetochores where it contributes to microtubule attachment and spindle formation.

At the end of G2, the cell has grown to a critical size and is prepared for M phase (Fig. 1g). The NE breaks down (NEBD) and NPCs are disassembled (Fig. 1h). Whereas most Nups disperse throughout the cytoplasm and the endoplasmic reticulum (ER)^16,17^, ELYS, together with components of the Nup107-160 complex and RANBP2, play an important role in prophase by binding to the kinetochores of recently replicated sister chromatids where they contribute to microtubule recruitment and spindle assembly (Fig. 1i)^18–20^. At telophase, when the chromatids are fully separated, the spindle and kinetochores dissolve and ELYS becomes tethered to thousands of sites on chromatin through its C-terminal chromatin binding region where it promotes chromatin decondensation^15,21^. In G1, the N-terminal half of ELYS provides a scaffold to bind the Nup107-160 complex and other ER-associated Nups, thereby coupling the initiation of NPC assembly with euchromatin formation, restoration of nucleocytoplasmic transport and transcription^11,22,23^ and completing the cycle depicted in Fig. 1.

Given this diversity of functions throughout the cell cycle, it is not surprising that ELYS is an Achilles’ heel of highly proliferative cells^8,9^, and this prompted us to ask whether this vulnerability extrapolates to cancer cells *in vivo*. To do this, we employed a highly tractable zebrafish model of hepatocellular carcinoma (HCC) in which hepatocyte hyperplasia and liver enlargement are driven by an inducible, hepatocyte-specific *EGFP*-*kras^G12V^* transgene^24^. In this study, we used two-photon microscopy to accurately quantitate liver volume in the presence and absence of a single nonsense mutation in one allele of the *ahctf1* locus^8,9^. Remarkably, we observed a pronounced reduction in tumour burden in response to this mild (50%) reduction in *ahctf1* gene dosage, while the rest of the animal was unaffected. We further show that this selective vulnerability of oncogenic *kras^G12V^*-expressing liver cancer cells to *ahctf1* heterozygosity possesses the hallmarks of a synthetic lethal interaction that may be exploited therapeutically.

## RESULTS

### *ahctf1* heterozygosity reduces tumour burden in a zebrafish *kras^G12V^-*driven model of hepatocellular carcinoma

In the zebrafish *kras^G12V^*-driven HCC model, the doxycycline-induced expression of a single *EGFP-kras^G12V^* transgene (genotype denoted *TO(kras^G12V^)^T/+^*) between 2-7 days post-fertilisation (dpf) leads to the accumulation of a constitutively active, EGFP-tagged oncogenic form of Kras specifically in hepatocytes^24^ (Fig. 2a). This forced expression of *EGFP-kras^G12V^* causes hepatocyte hyperplasia and a substantial increase in liver volume that can be quantified by two-photon microscopy. To investigate the requirement for Elys in this cancer setting, we introduced a mutant *ahctf1* allele (containing a nonsense mutation at codon 1319) from a zebrafish development mutant (*flo*^ti262^)^7^ into the genome of this model. This resulted in a 57% reduction in *ahctf1* mRNA expression in *ahctf1^+/−^* larvae at 7 dpf (Fig. 2b), indicating that the premature codon encoded by the nonsense mutation triggers nonsense mediated decay rather than expression of a truncated Elys protein.

**Fig. 2.**
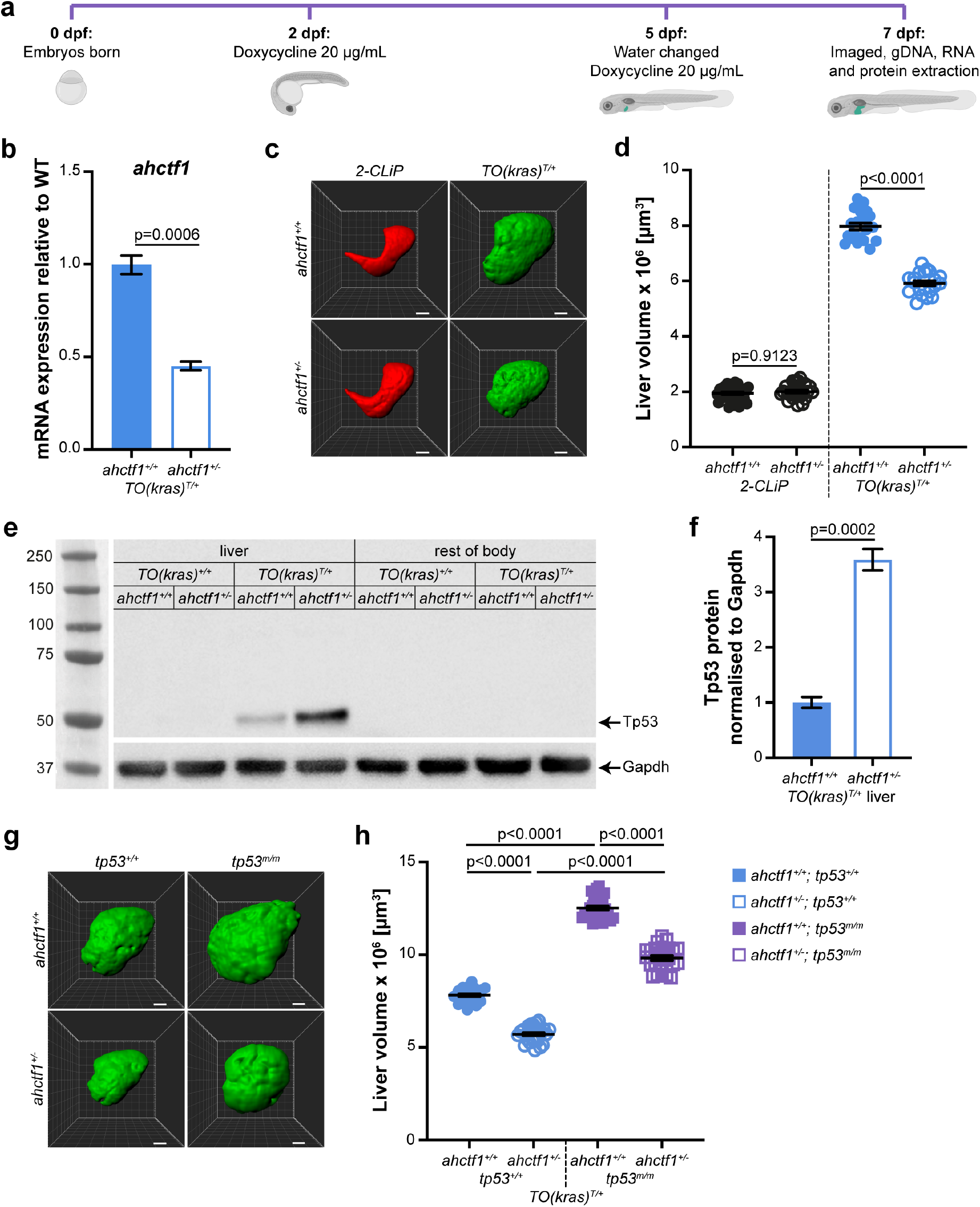
*ahctf1* heterozygosity reduces liver volume in a zebrafish *kras^G12V^-*driven model of hepatocellular carcinoma in both Tp53 proficient and deficient larvae. **a.** Protocol for doxycycline induction of *TO(kras^G12V^)^T/+^* expression in developing zebrafish larvae. **b.** RT-qPCR analysis of *ahctf1* mRNA levels in pooled micro-dissected livers (n=3 biological replicates). **c.** Representative three-dimensional reconstructions of *2-CLiP* and *TO(kras^G12V^)^T/+^* livers of the indicated *ahctf1* genotype. Scale bar 25 µm. **d.** Impact of *ahctf1* heterozygosity on liver volume in *2-CLiP* and *TO(kras^G12V^)^T/+^* larvae (n≥20). **e.** Representative western blot of Tp53 protein signals in lysates of *TO(kras^G12V^)^T/+^* larvae of the indicated *ahctf1* genotype. **f.** Quantification of Tp53 protein levels normalised by reference to the Gapdh loading control (n=3 independent experiments). **g.** Representative three-dimensional reconstructions of *TO(kras^G12V^)^T/+^* livers of the indicated *ahctf1* and *tp53* genotypes. Scale bar 25 µm. **h.** Impact of *ahctf1* heterozygosity and homozygous *tp53* mutation on liver volume in *2-CLiP* and *TO(kras^G12V^)^T/+^* larvae (n≥20). Data are expressed as mean ± SEM. Significance was calculated using a Student’s t-test or one-way ANOVA with Tukey’s multiple comparisons test.

To determine the effect of *ahctf1* heterozygosity (and other treatments) on normal liver cells, we used a control transgenic line, denoted *2-CLiP* (2-<u>C</u>olour <u>Li</u>ver <u>P</u>ancreas), in which hepatocytes express dsRed fluorescence but no oncogenic transgene^25^. Mean liver volume in this model at 7 dpf was 1.95 x 10^6^ ± 4.99 x 10^4^ μm^3^ and did not change in response to *ahctf1* heterozygosity (Fig. 2c,d), consistent with previous observations that *ahctf1* heterozygotes develop normally, reach sexual maturity, and exhibit a normal lifespan^9^. In the *TO(kras^G12V^)^T/+^* model, induced expression of oncogenic Kras^G12V^ leads to hepatocyte hyperplasia and a striking (4.1-fold) increase in liver volume to 7.97 x 10^6^ ± 1.21 x 10^5^ μm^3^, an increment of 6.02 x 10^6^ μm^3^ over the course of 5 d of dox treatment. Remarkably, the potency of this strong oncogenic *kras^G12V^* signal was pared back by *ahctf1* heterozygosity (down to 5.92 x 10^6^ ± 8.83 x 10^4^ µm^3^), equating to a 35% reduction in excess liver volume. These data demonstrate that a 50% reduction in *ahctf1* gene dosage markedly restricts tumour burden in livers expressing oncogenic *kras^G12V^*, while having no effect on normal livers. Thus, mildly disrupted *ahctf1* expression is a selective vulnerability of cancer cells.

Robust and persistent overexpression of RAS oncoproteins is frequently associated with oncogene-induced stress comprising defects in DNA replication, DNA damage and genome instability. In the presence of wildtype TP53, this damage is limited by activation of TP53 transcription pathways that induce cell cycle arrest, senescence and apoptosis. To determine whether expression of *kras^G12V^* causes oncogene-induced stress and Tp53 activation in our model, we measured the levels of Tp53 protein in pooled lysates of micro-dissected *TO(kras^G12V^)^+/+^* and *TO(kras^G12V^)^T/+^* livers and the remains of the larvae (Fig. 2e,f). No Tp53 signal was obtained from non-*kras^G12V^*-expressing livers or the body remains after liver removal. We detected a weak Tp53 signal in extracts of *ahctf1^+/+^;TO(kras^G12V^)^T/+^* livers, indicating that expression of the *kras^G12V^* oncogene alone elicited a weak Tp53 response. In contrast, a much stronger Tp53 signal (>3.5-fold) was detected in *ahctf1^+/−^;TO(kras^G12V^)^T/+^* livers, demonstrating a more severe level of stress in heterozygous *ahctf1* hepatocytes expressing oncogenic *kras^G12V^* than occurs in hepatocytes that are WT for *ahctf1*.

### Loss of Tp53 function enhances *kras^G12V^*-driven liver enlargement, which is partially alleviated by *ahctf1* heterozygosity

To test whether activation of the Tp53 pathway was involved in reducing the volume of *kras^G12V^*-expressing livers on a heterozygous *ahctf1* background, we repeated the experiment in the absence of Tp53 function. Here, we utilised the zebrafish *tp53^e7^* or *tp53^M214K^* allele, which changes a conserved amino acid residue within the Tp53 DNA-binding domain corresponding to a mutational hotspot in human cancer, producing a transactivation-dead Tp53 variant^26^. Abrogating Tp53 function by homozygous expression of this allele (denoted *tp53^m/m^*) in *ahctf1^+/+^*;*TO(kras^G12V^)^T/+^* larvae supported a further large increase (82%) in liver volume (1.25 x 10^7^ ± 1.10 x 10^5^ μm^3^) compared to livers on the *tp53^+/+^* background (Fig. 2g,h) (Supplementary Fig. 1-3), demonstrating that Tp53 function normally restrains tumour growth and liver enlargement in this model. Interestingly, the increase in liver volume was significantly blunted (35% reduction) on a heterozygous *ahctf1* background, despite the absence of functional Tp53.

### *ahctf1* heterozygosity triggers cell death in *TO(kras^G12V^)^T/+^* hepatocytes through Tp53-dependent and independent mechanisms

We previously showed that Tp53 protein was upregulated in the livers of *(kras^G12V^)^T/+^* larvae on a heterozygous *ahctf1* background (Fig. 2e). To determine whether this activated a Tp53-dependent cell death response, we utilised a transgenic line, Tg(*actb2:SEC-Hsa.ANXA5-mKate2,cryaa:mCherry*)^uq24rp^ (hereafter denoted *Annexin 5-mKate*) that constitutively expresses a fusion protein comprising Annexin 5 and the far-red fluorophore mKate^2^^7^. This fusion protein gives rise to discrete fluorescent puncta in cells undergoing apoptosis by binding to phosphatidylserine that is normally inaccessible on the inner leaflet of the plasma membrane but is exposed as the membrane breaks down.

Annexin 5 fluorescent puncta were sparsely distributed (1.5 puncta per 10^−5^ μm^3^) throughout the livers of *ahctf1^+/+^;tp53^+/+^;TO(kras^G12V^)^T/+^* larvae demonstrating that expression of *kras^G12V^* alone is associated with low levels of cell death (Fig. 3a,b). However, the number of Annexin 5 fluorescent objects was 3.5-fold higher (5.2 puncta per 10^−5^ μm^3^) in the livers of *ahctf1^+/−^;tp53^+/+^;TO(kras^G12V^)^T/+^* larvae demonstrating that the reduction in tumour burden caused by *ahctf1* heterozygosity is due, at least in part, to an increase in cell death (Fig. 3a,b left two columns). These results were recapitulated when the cleaved, (active form) of caspase 3 was used as an alternative marker of apoptosis (Supplementary Fig. 4). That cell death in hyperplastic hepatocytes is largely dependent on Tp53 function was demonstrated by negligible apoptosis in the livers of *ahctf1^+/+^;tp53^m/m^;TO(kras^G12V^)^T/+^* larvae (Fig. 3a,b right two columns). When this experiment was repeated on a heterozygous *ahctf1^+/−^* background, we observed a 17.6-fold increase in the abundance of Annexin 5 fluorescent puncta compared to livers on a WT *ahctf1* background. However, overall, there were significantly fewer Annexin 5 puncta in heterozygous *ahctf1* hepatocytes on *tp53^m/m^* background compared to those expressing WT *tp53*. Collectively, these data identify a novel synthetic lethal interaction between *kras^G12V^* and one mutated allele of *ahctf1* that is mediated, at least in part, by Tp53.

**Fig. 3.**
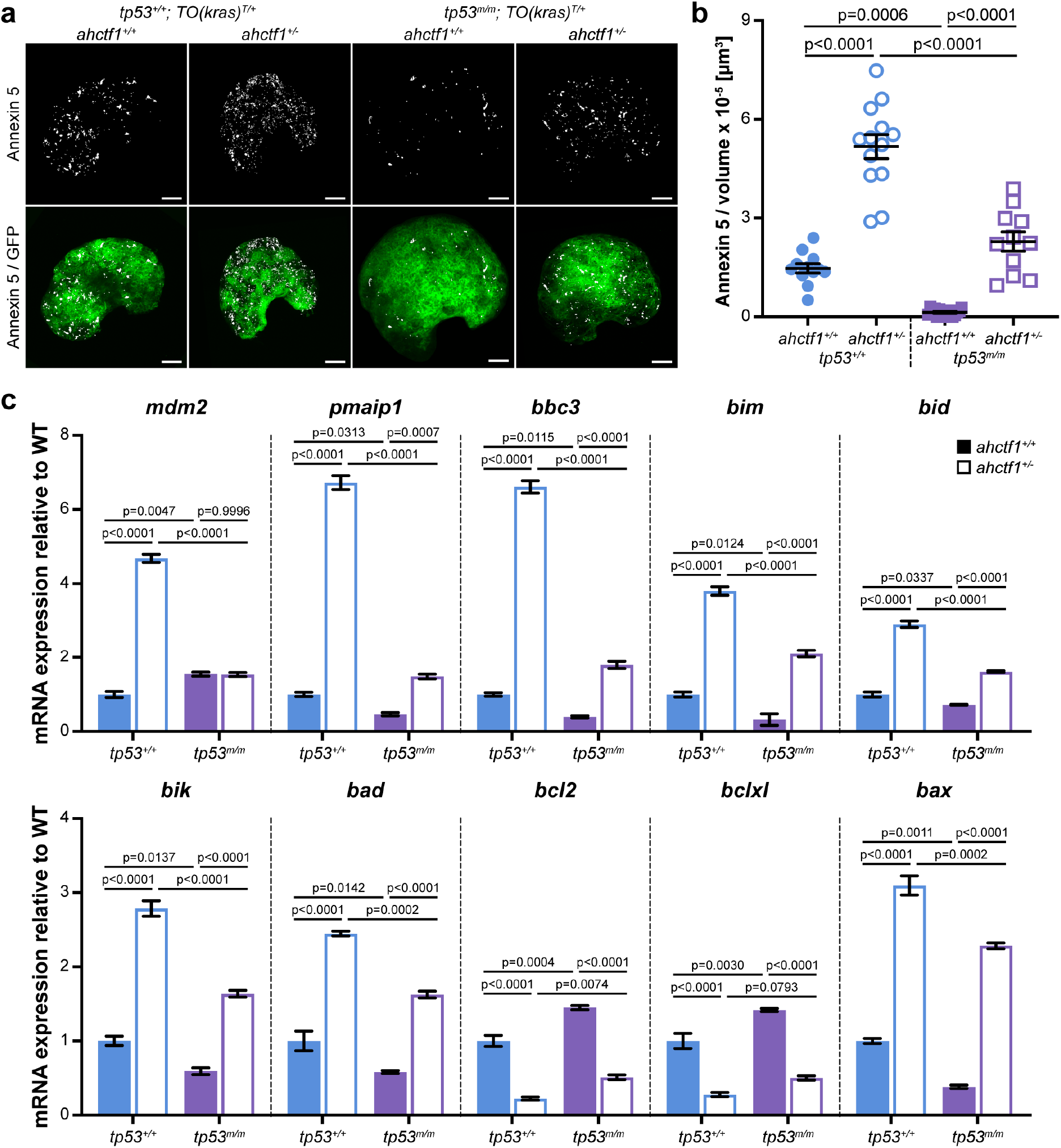
*ahctf1* heterozygosity increases cell death in *TO(kras^G12V^)^T/+^* hepatocytes in both Tp53 proficient and deficient larvae. **a.** Representative maximum intensity projection images of Annexin 5-mKate fluorescence (white puncta), indicating cells undergoing apoptosis in *TO(kras^G12V^)^T/+^* livers of the indicated *ahctf1* and *tp53* genotypes. Scale bar 25 µm. **b.** Quantification of the density of Annexin 5 fluorescent foci in *TO(kras^G12V^)^T/+^* livers of the indicated *ahctf1* and *tp53* genotypes (n≥11). **c.** RT-qPCR analysis of the specified mRNAs in *TO(kras^G12V^)^T/+^* micro-dissected livers of the indicated *ahctf1* and *tp53* genotypes (n=3 biological replicates). Data are expressed as mean ± SEM. Significance was calculated using a one-way ANOVA with Tukey’s multiple comparisons test.

We showed that the elevated level of Tp53 protein observed in *ahctf1^+/−^;TO(kras^G12V^)^T/+^* livers, compared to *ahctf1^+/+^;TO(kras^G12V^)^T/+^* livers (Fig. 2e,f), was transcriptionally active by the upregulated (4.7-fold) mRNA expression of a canonical Tp53 target gene, *mdm2* (Fig. 3c). This prompted us to quantify the mRNA expression levels of a suite of Bcl2 family genes that regulate intrinsic (mitochondrial) apoptosis (Fig. 3c). Direct Tp53 transcriptional targets, *pmaip1* (aka *noxa*) and *bbc3* (aka *puma*), were both upregulated by >6.5-fold in *ahctf1^+/−^;tp53^+/+^;TO(kras^G12V^)^T/+^* livers compared to *ahctf1^+/+^;tp53^+/+^;TO(kras^G12V^)^T/+^* livers. mRNAs encoding the other BH3-only apoptosis effector proteins, Bim, Bid, Bik and Bad, and the pro-apoptotic executioner protein, Bax were also increased in *ahctf1^+/−^;tp53^+/+^;TO(kras^G12V^)^T/+^* livers compared to *ahctf1^+/+^;tp53^+/+^;TO(kras^G12V^)^T/+^* livers. Meanwhile, transcripts encoding Bcl2 and Bclxl pro-survival proteins were significantly downregulated by *ahctf1* heterozygosity. In the absence of Tp53 function, *ahctf1* heterozygosity still increased the expression of these pro-apoptotic transcripts, albeit less markedly. All these data are consistent with induction of Tp53 target genes playing a major role in the cell death response to *ahctf1* heterozygosity in *kras^G12V^*-expressing livers. Notwithstanding this, we observed some residual cell death concomitant with reduced tumour burden in *ahctf1* heterozygotes, even in the absence of Tp53 function.

### *ahctf1* heterozygosity restricts DNA replication in *TO(kras^G12V^)^T/+^* hepatocytes

To explore whether the reduction in liver enlargement we observe in *ahctf1^+/−^*;*TO(kras^G12V^)^T/+^* larvae also involves impaired cell cycle progression, we used an EdU incorporation assay to identify cells in S phase (Fig. 4a). We observed 32% EdU-positive cells in the enlarged livers of *ahctf1^+/+^;tp53^+/+^;TO(kras^G12V^)^T/+^* larvae and >20% EdU-positive cells (32% reduction) in the livers of *ahctf1^+/−^; tp53^+/+^;TO(kras^G12V^)^T/+^* larvae (Fig. 4b,c). In Tp53 deficient larvae, liver volume was markedly increased as expected (Fig. 4b) and there was a doubling in the percentage of EdU-positive cells (63% compared to 32%) (Fig. 4c). *ahctf1* heterozygosity reduced liver volume in both Tp53 proficient and deficient larvae (Fig. 4b), consistent with Fig. 2, and reduced the percentage of EdU-positive cells in both Tp53 proficient (22% compared to 32%) and Tp53 deficient larvae (53% compared to 63%) (Fig. 4c). These data indicate that the overall reduction in liver tumour burden we observe in response to *ahctf1* heterozygosity is due to a combination of fewer cells undergoing DNA replication and more cells undergoing apoptosis.

**Fig. 4.**
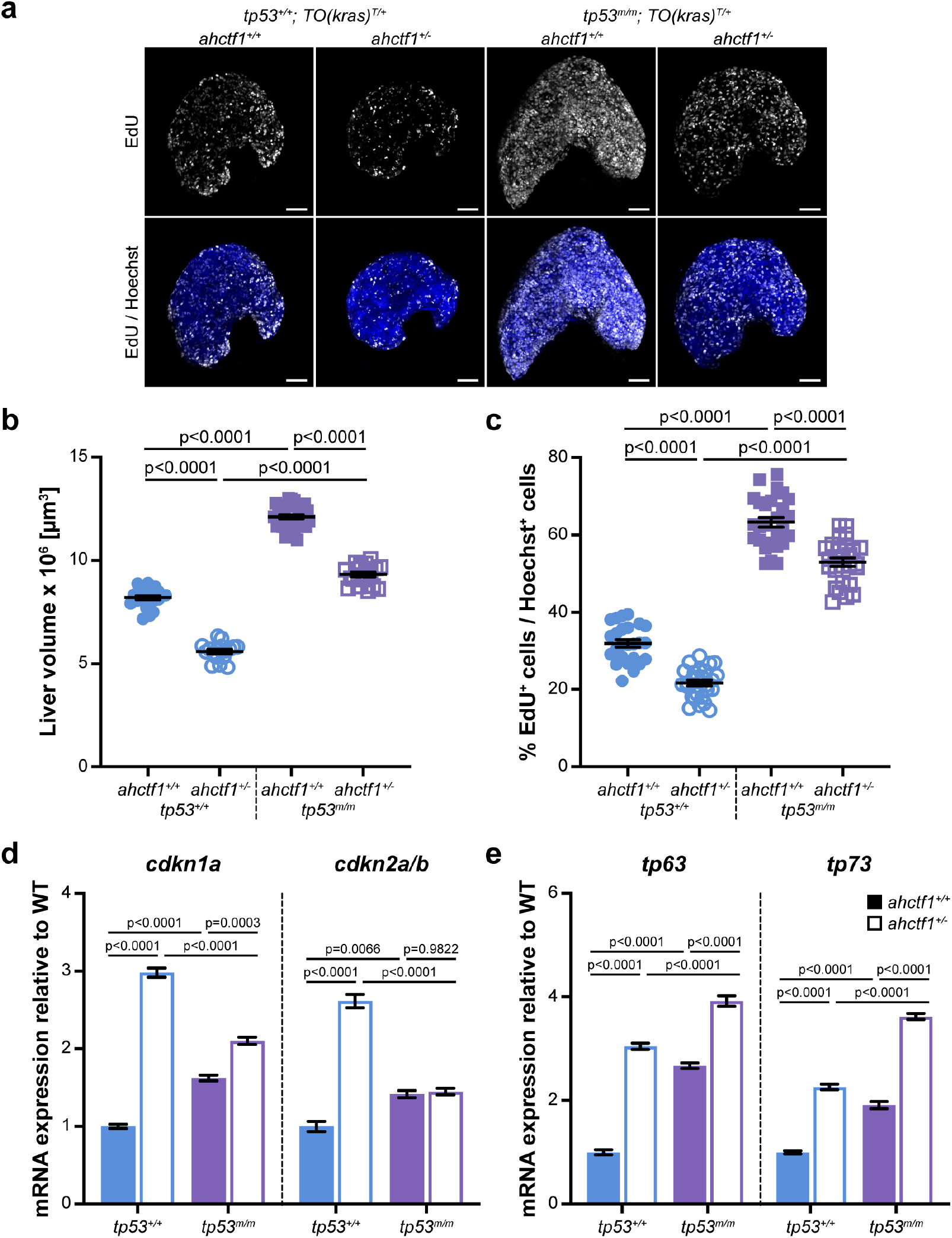
*ahctf1* heterozygosity restricts DNA replication in *TO(kras^G12V^)^T/+^* hepatocytes. **a.** Representative maximum intensity projection images of EdU incorporation (white puncta) into *TO(kras^G12V^)^T/+^* livers of the indicated *ahctf1* and *tp53* genotypes. Scale bar 25 µm. **b.** Impact of *ahctf1* heterozygosity and homozygous *tp53* mutation on liver volume in *2-CLiP* and *TO(kras^G12V^)^T/+^* larvae (n≥17). **c.** Quantification of the percentage of EdU positive nuclei per Hoechst 33342 positive nuclei (n≥25). **d.** RT-qPCR analysis of mRNA expression of the cell cycle regulators, *cdkn1a* and *cdkn2a/b* and **e.** *tp63* and *tp73* in *TO(kras^G12V^)^T/+^* micro-dissected livers of the indicated *ahctf1* and *tp53* genotypes (n=3 biological replicates). Data are expressed as mean ± SEM. Significance was assessed using a one-way ANOVA with Tukey’s multiple comparisons test.

To investigate the molecular mechanisms underlying this observation, we performed RT-qPCR analysis of the negative cell cycle regulators, *cdkn1a* and *cdkn2a/b* in the presence and absence of Tp53. We found that *cdkn1a* and *cdkn2a/b*, encoding p21 and p16^Ink4a^ respectively, were upregulated in *ahctf1^+/−^ ;tp53^+/+^*;*TO(kras^G12V^)^T/+^* livers compared to *ahctf1^+/+^;tp53^+/+^*;*TO(kras^G12V^)^T/+^* livers (Fig. 4d). For *cdkn1a*, this response to *ahctf1* heterozygosity was partially dependent on intact Tp53 function. In contrast, the level of *cdkn2a/b* mRNA expression was increased in response to *ahctf1* heterozygosity in *tp53^+/+^;TO(kras^G12V^)^T/+^* livers, but there was no response to *ahctf1* heterozygosity in *tp53^m/m^* livers, consistent with *cdkn2a/b* expression being regulated solely by Tp53.

To determine how *ahctf1* heterozygosity reduces tumour burden in the absence of Tp53, we investigated the impact of *ahctf1* heterozygosity on the expression of *tp53* family members, *tp63* and *tp73*. *tp63* and *tp73* mRNA expression levels were up-regulated by 3.0-fold and 2.3-fold, respectively, in heterozygous *ahctf1*, *tp53^+/+^;TO(kras^G12V^)^T/+^* livers, compared to WT *ahctf1* livers. While *tp63* and *tp73* transcripts were upregulated in the absence of *tp53* (*tp53^m/m^*) for both *ahctf*1 genotypes, their levels were significantly higher in *ahctf1* heterozygotes. Taken together, the data in Figs. 3 and 4 suggest that *ahctf1* heterozygosity constrains *kras^G12V^*-driven hepatocyte hyperplasia and liver enlargement by activating Tp53 transcriptional programs that induce cell cycle arrest and cell death. Furthermore, the beneficial effect of *ahctf1* heterozygosity to reduce tumour burden persists in the absence of functional Tp53, due to compensatory increases in the levels of *tp63* and *tp73*.

### *ahctf1* heterozygosity disrupts the abundance of nuclear pore complexes in *TO(kras^G12V^)^T/+^* hepatocytes

To determine how *ahctf1* heterozygosity leads to Tp53 activation, we examined the integrity of some of the essential processes in which the Elys protein plays a role. As depicted in Fig. 1, Elys fulfils many distinct functions during the cell cycle. Starting with post-mitotic nuclear pore assembly (Fig. 1a), we used Airyscan confocal laser-scanning microscopy to look at the abundance and distribution of NPCs in thick sections of liver (200 μm) stained with an antibody (mAb414) that recognises FG-repeat Nups (Nup358, 214, 153 and 62) in mature NPCs. Non-*kras^G12V^* expressing hepatocytes exhibited fluorescent puncta corresponding to NPCs at the nuclear rim with negligible staining in the cytoplasm, a pattern that was unaffected by *ahctf1* genotype (Fig. 5a; left two columns). Upon induction of expression of the *TO(kras^G12V^)^T/+^* transgene, fluorescence intensity was markedly increased in the presence of wildtype *ahctf1* (Fig 5a, third column) and there was an increase in the ratio of nuclear:cytoplasmic staining (Fig 5b). However, fluorescence intensity at the nuclear rim was diminished in *ahctf1* heterozygotes, concomitant with the appearance of puncta in the cytoplasm (Fig. 5a, fourth column; arrows), leading to a 21% reduction in the ratio of nuclear:cytoplasmic fluorescence intensity (Fig. 5b).

**Fig. 5.**
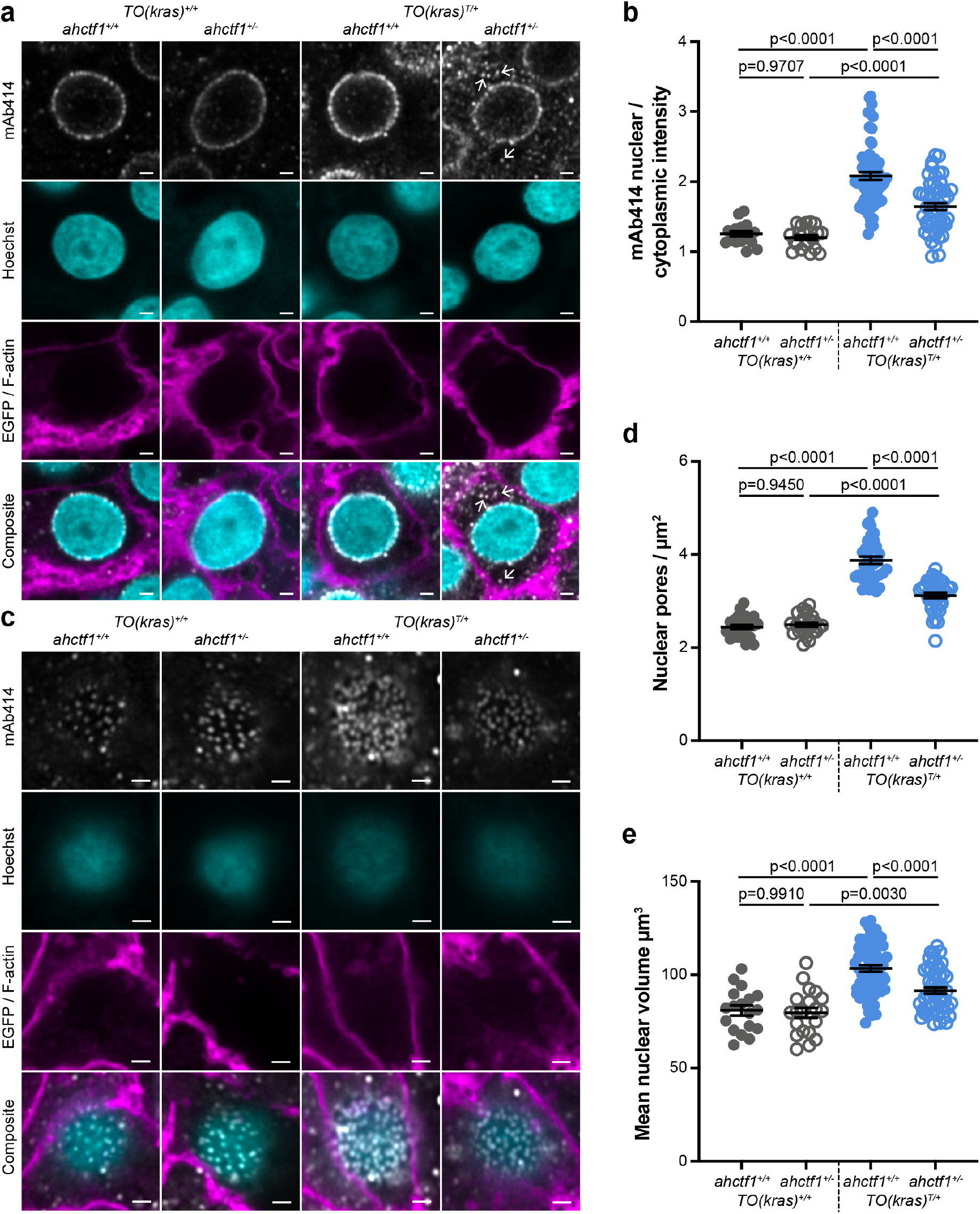
*ahctf1* heterozygosity disrupts nuclear pore complexes in *TO(kras^G12V^)^T/+^* hepatocytes. **a.** Representative Airyscan imaging of liver sections stained with mAb414 (white) marking FG-Nups, Hoechst 33342 (cyan) marking DNA, and rhodamine phalloidin (magenta) marking the F-actin cytoskeleton in non-*TO(kras^G12V^)*-expressing cells or EGFP-Kras^G12V^ (magenta) marking the cell membrane in *TO(kras^G12V^)*-expressing cells of the indicated *ahctf1* and *TO(kras^G12V^)* genotypes. Arrows in right hand panel point to mAb414/FG-nucleoporin staining in the cytoplasm. Scale bar 2 µm. **b.** Quantification of mean nuclear/cytoplasmic fluorescence intensity of mAb414 staining after 3D segmentation and morphological filtering of nuclear and cytoplasmic areas (n>18). **c.** Representative Airyscan images of mAb414 staining at the nuclear surface of sections of the indicated *ahctf1* and *TO(kras^G12V^)* genotype. Scale bar 1 µm**. d.** Quantification of nuclear pore density (n≥25). **e.** Quantification of nuclear volume (n≥25). Data are expressed as mean ± SEM. Significance was assessed using a one-way ANOVA with Tukey’s multiple comparisons test.

To determine the abundance of NPCs, we analysed images acquired at the nuclear surface (Fig. 5c). The pattern and density of fluorescent puncta observed at the nuclear surface of non-*kras^G12V^* expressing hepatocytes was not affected by *ahctf1* genotype (Fig. 5c; left two columns). The expression of the *kras^G12V^* transgene in the presence of wildtype *ahctf1* resulted in 59% more fluorescent puncta/NPCs at the nuclear surface of hyperplastic hepatocytes, which was reduced by 20% in response to *ahctf1* heterozygosity (Fig. 5c, d; right two columns). Induced *kras^G12V^* expression also produced a 28% increase in nuclear volume compared to non-*kras^G12V^* expressing cells (Fig. 5e), and this increase was reduced to 13% by *ahctf1* heterozygosity. Together, these data indicate that hyperplastic hepatocytes expressing the *kras^G12V^* oncogene contain more abundant nuclear pores and larger nuclei than their non-*kras^G12V^* expressing counterparts, properties that are likely to enhance the trafficking of mRNAs and proteins in and out of the nucleus to support the transcriptional and translational needs of rapidly cycling cancer cells. However, *ahctf1* heterozygosity reduced NPC density in *kras^G12V^*-hepatocytes, potentially restricting the efficiency of nucleocytoplasmic trafficking.

### *ahctf1* heterozygosity impairs mitotic spindle assembly and chromosome segregation in *TO(kras^G12V^)^T/+^* hepatocytes

Next, we examined the impact of reduced Elys expression on spindle formation and chromosome segregation (Fig. 1b). We assessed these features in cryosections of liver using *α*-tubulin and DAPI to stain microtubules and chromatin, respectively. Metaphase cells in *ahctf1^+/+^;TO(kras^G12V^)^T/+^* livers exhibited normal bipolar spindle formation followed by complete chromosome segregation during anaphase (Fig. 6a). In contrast, metaphase cells in *ahctf1^+/−^;TO(kras^G12V^)^T/+^* hepatocytes displayed abnormal multipolar spindles and misaligned chromosomes (Fig. 6b). Proper chromosome segregation was disrupted with multiple anaphase bridges formed. While the distribution of cells observed at different mitotic stages was similar in *ahctf1^+/+^* and *ahctf1^+/−^* hepatocytes (Fig. 6c), mitotic abnormalities were observed in 50% of *ahctf1^+/−^;TO(kras^G12V^)^T/+^* hepatocytes during metaphase and anaphase but not at all in *ahctf1^+/+^;TO(kras^G12V^)^T/+^* hepatocytes (Fig. 6d). Taken together, these data suggest that hyperplastic hepatocytes require a full complement of *ahctf1* expression to maintain proper cell division.

**Fig. 6.**
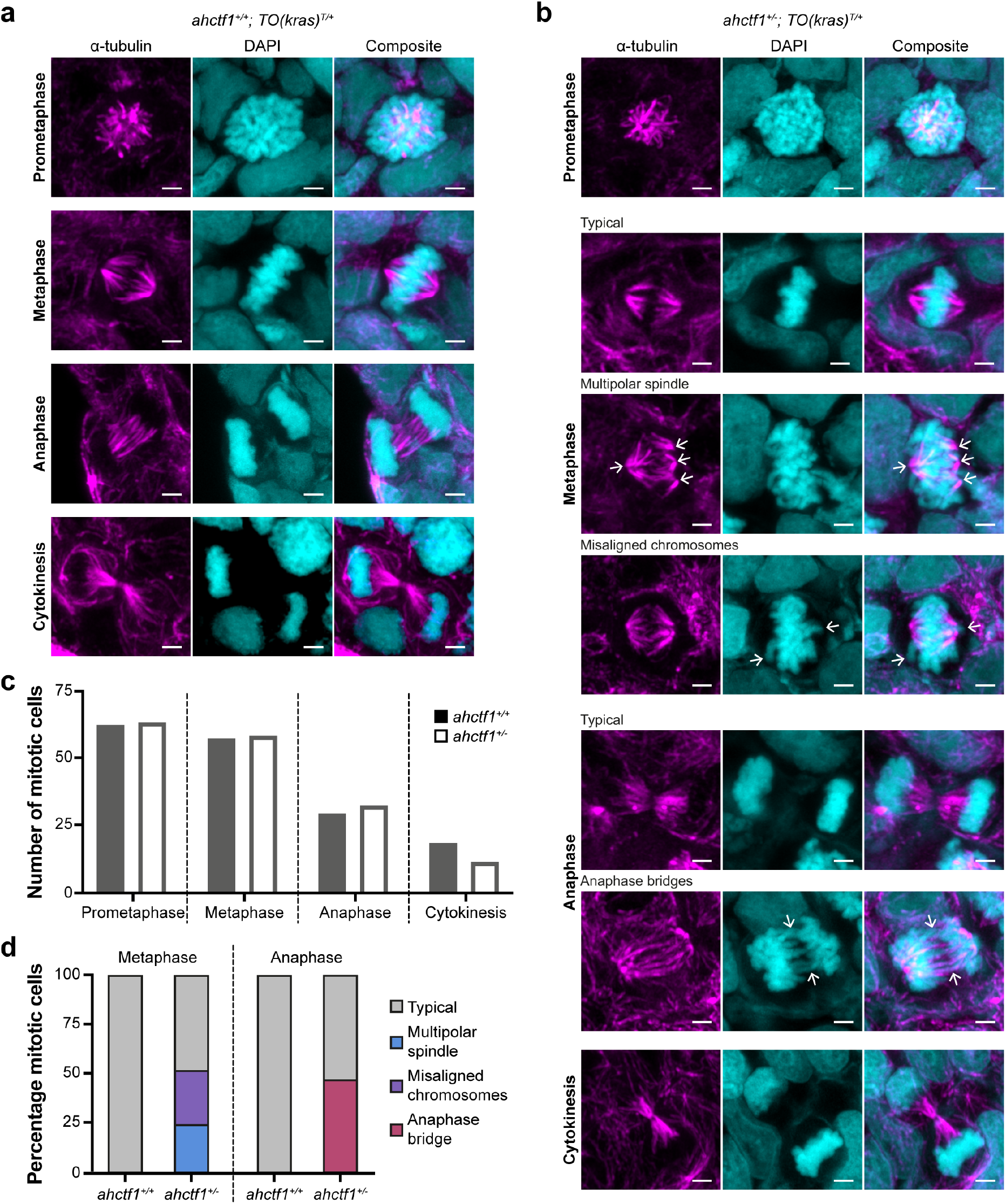
*ahctf1* heterozygosity impairs mitotic spindle assembly and chromosome segregation in *TO(kras^G12V^)^T/+^* hepatocytes. **a.** Representative Airyscan imaging of liver cryosections stained with *α*-tubulin antibody (magenta) marking spindle microtubules and DAPI (cyan) marking DNA in mitotic cells of *TO(kras^G12V^)^T/+^* larvae on a wildtype *ahctf1^+/+^* background. **b.** Mitotic cells in liver cryosections of *TO(kras^G12V^)^T/+^* larvae on a heterozygous *ahctf1^+/−^*background exhibit multiple defects, including multipolar spindles, misaligned chromosomes and anaphase bridges (arrows). Scale bar 2 µm. **c.** Distribution of cells observed at different mitotic stages (n=92 livers, 326 mitotic cells) **d.** Quantification of the percentage of mitotic hepatocytes exhibiting an aberrant phenotype (n=14-57).

### *ahctf1* heterozygosity causes DNA damage in *TO(kras^G12V^)^T/+^* hepatocytes

So far, our results demonstrate that a 50% decrease in *ahctf1* expression has multiple impacts in *TO(kras^G12V^)^T/+^* hepatocytes, including disrupted NPC assembly, impaired mitotic spindle assembly and incomplete chromosome segregation, consistent with oncogene-induced stress. As several of these defects can converge to produce DNA damage, we looked for evidence of DNA damage by staining cryosections of liver with DAPI and γ-H2AX to mark DNA double-stranded breaks (Fig. 7a,b). We found that only 1% of *ahctf1^+/+^;TO(kras^G12V^)^T/+^* hepatocyte nuclei were positive for γ-H2AX (Fig. 7c). This was increased by 6-fold in *ahctf1^+/−^;TO(kras^G12V^)^T/+^* hepatocyte nuclei.

**Fig. 7.**
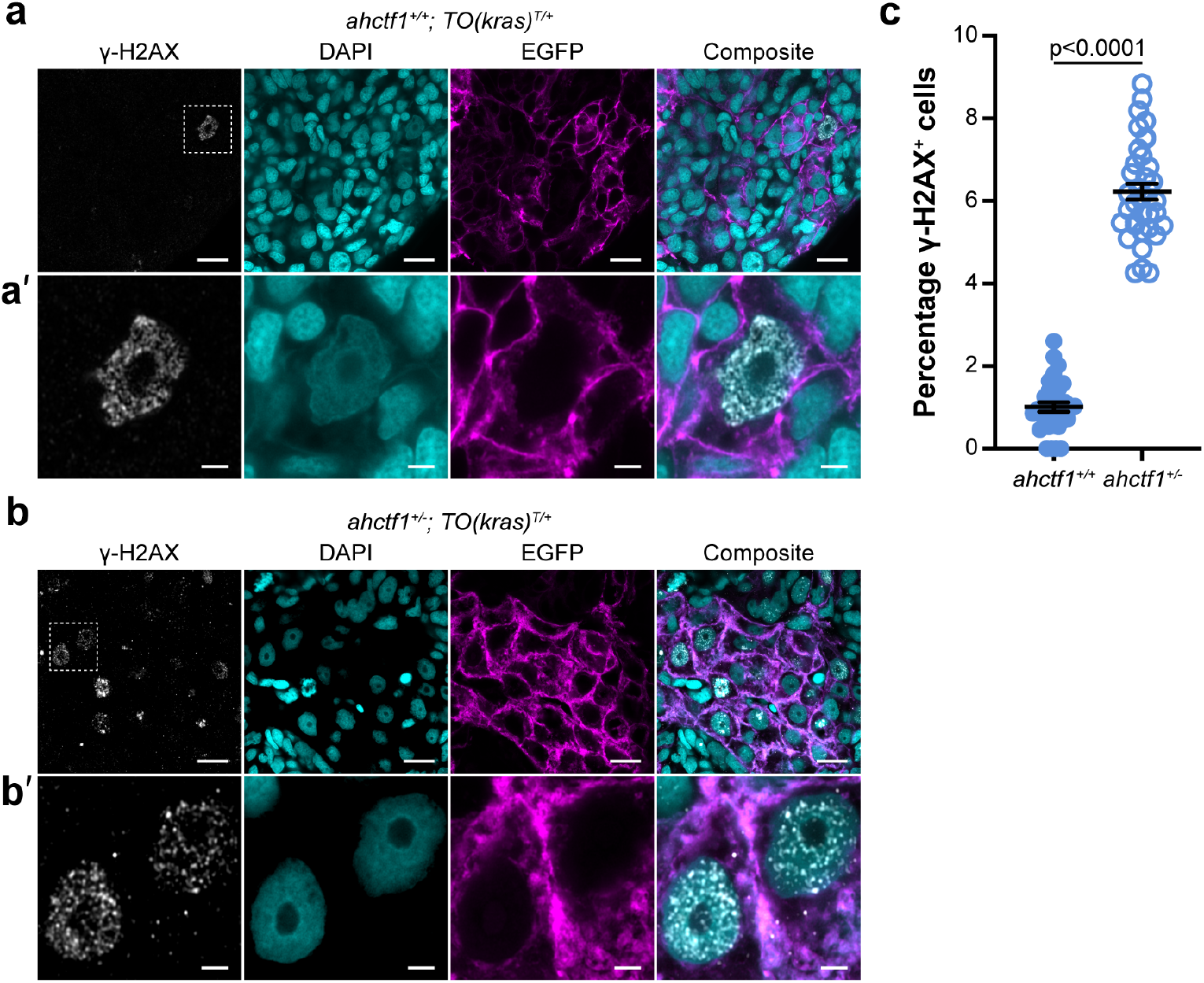
*ahctf1* heterozygosity leads to accumulation of DNA double-strand breaks in *TO(kras^G12V^)^T/+^* hepatocytes. **a.** Representative Airyscan imaging of cryosections of liver from *ahctf1^+/+^;TO(kras^G12V^)^T/+^* larvae stained with γ-H2AX antibody (white) marking DNA double-strand breaks, DAPI (cyan) marking DNA and EGFP-Kras^G12V^ (magenta) marking the cell membrane. Scale bar 5 µm. **a’.** Inset of γ-H2AX positive nuclei in *ahctf1^+/+^;TO(kras^G12V^)^T/+^* hepatocytes. Scale bar 2 µm. **b.** Representative images of cryosections of liver from *ahctf1^+/−^;TO(kras^G12V^)^T/+^* larvae. Scale bar 5 µm. **b’.** Inset of γ-H2AX positive nuclei in cryosections of liver from *ahctf1^+/−^;TO(kras^G12V^)^T/+^* larvae. Scale bar 2 µm. **c.** Quantification of the percentage of hepatocytes positive for γ-H2AX (n≥31). Data are expressed as mean ± SEM. Significance was calculated using a Student’s t-test.

Collectively, these data show that the proper functioning of diverse cellular processes in hyperplastic hepatocytes depends on wildtype levels of *ahctf1* expression. We show that heterozygous expression of *ahctf1* in mutant Kras-fuelled cancer cells leads to multiple insufficiencies that combine to amplify oncogene-induced stress and cause DNA damage, in so doing providing a stimulus to activate Tp53 and transcriptional programs that trigger cell cycle arrest and apoptosis.

### Combinatorial approaches can completely restrict *kras^G12V^*-driven liver hyperplasia

Having determined that heterozygous *ahctf1* mutation is effective in restricting *kras^G12V^*-driven hepatocyte hyperplasia and liver enlargement, we investigated the potential of targeting cancer via another nucleoporin, Ranbp2, in our zebrafish model of HCC. RANBP2 (also known as NUP358) is a major component of the cytoplasmic filaments of NPCs, where it functions in numerous transport pathways. Like ELYS, RANBP2 also plays non-canonical roles outside of the NPC, including in mitotic progression and the maintenance of genome integrity^28–30,31^.

We identified our zebrafish *ranbp2* mutant in a transgene assisted ENU-mutagenesis screen for mutations affecting endodermal organ morphogenesis^32^ and showed, using whole genome sequencing and homozygosity mapping^33,34^, that the underlying mutation was a nonsense mutation in *ranbp2* (Supplementary Fig. 5). Like several other mutants we identified in this screen^8,35,36^, mutant *ranbp2* larvae exhibit morphological deficiencies in proliferative compartments, including the developing intestinal epithelium, craniofacial complex and eye (Supplementary Fig. 5).

We found that *ranbp2* heterozygosity alone or in combination with *ahctf1* heterozygosity did not impact on liver volume in *2-CLiP* larvae (Fig. 8a,b). In *TO(kras^G12V^)^T/+^* larvae, liver volume was increased 3.8-fold, as observed previously, and the excess liver volume was reduced by 38% and 13% by the introduction of a single mutation in *ahctf1* and *ranbp2*, respectively (Fig. 8b). Remarkably, we found a striking synergistic benefit of trans heterozygosity in *ranbp2^+/−^;ahctf1^+/−^;TO(kras^G12V^)^T/+^* larvae, to the extent that liver volume was no longer distinguishable from that of normal *2-CLiP* livers. These results show that cancer cells are highly susceptible to combinatorial targeting of Nup function and that it may be possible to exploit this vulnerability to produce highly beneficial outcomes without impacting negatively on healthy tissues.

**Fig. 8.**
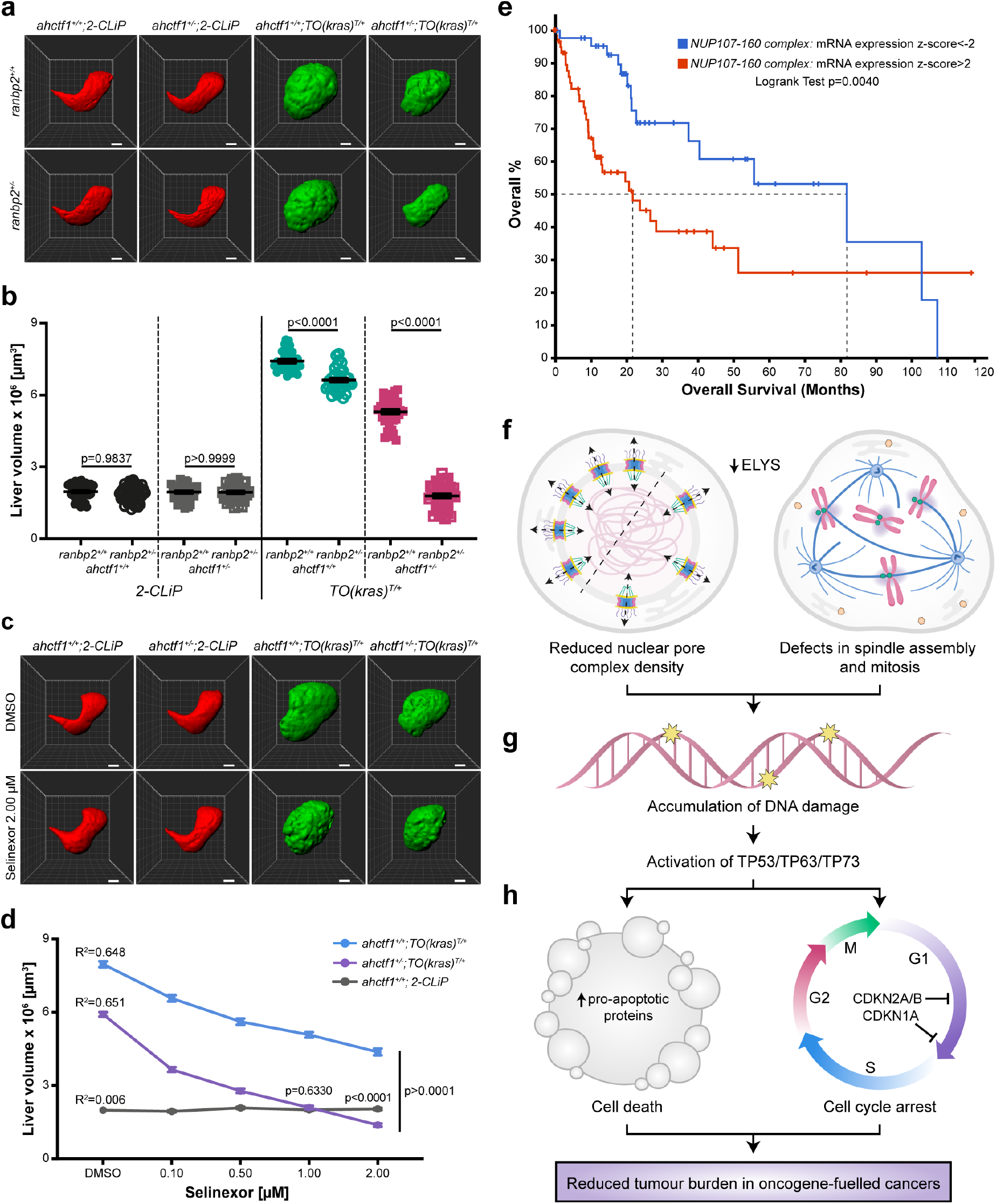
Combined approaches completely restrict *kras^G12V^*-driven liver hyperplasia. **a.** Representative three-dimensional reconstructions of livers from *2-CLiP* and *TO(kras^G12V^)^T/+^* larvae of the indicated *ahctf1* and *ranbp2* genotypes. Scale bar 25 µm. **b.** Impact of *ahctf1* heterozygosity and *ranbp2* heterozygosity on liver volume in *2-CLiP* and *TO(kras^G12V^)^T/+^* larvae (n≥30). Significance was calculated using a one-way ANOVA with Tukey’s multiple comparisons test. **c.** Representative three-dimensional reconstructions of *TO(kras^G12V^)^T/+^* livers of the indicated *ahctf1* genotype treated with DMSO or 2.00 µM Selinexor from 5-7 dpf. **d.** Dose-dependent impact of Selinexor treatment (0.10-2.00 µM) on liver volume in *2-CLiP* (grey), *ahctf1^+/+^;TO(kras^G12V^)^T/+^* (blue) and *ahctf1^+/−^;TO(kras^G12V^)^T/+^* (purple) larvae (n≥20). Data are expressed as mean ± SEM. Significance was calculated by linear regression analysis. **e.** Overall survival of HCC patients with mRNA expression z-scores >2 for one or more Nup107-160 complex components (red line; 63 cases); median overall survival 21.70 months. Overall survival of HCC patients with mRNA expression z-scores <-2 for one or more Nup107-160 complex components (blue line; 45 cases); median overall survival 81.73 months. Data from the TCGA LIHC dataset (total 372 samples) available in the cBioPortal for Cancer Genomics. **f.** Schematic depiction of two of the processes disrupted by mild depletion of Elys protein. **g.** Accumulation of DNA damage and activation of Tp53 transcription programs. **h**. Cell death and cell cycle arrest of hyperproliferative oncogene-expressing cancer cells.

To examine this concept in a therapeutic context, we assessed the effect of Selinexor, a selective inhibitor of nuclear export (SINE) in our *kras^G12V^*-driven liver hyperplasia model. Selinexor is an XPO1 (exportin) inhibitor with FDA approval for the treatment of refractory or relapsed multiple myeloma and diffuse large B-cell lymphoma and is in clinical trials for several other cancers, including HCC^37,38^. While we observed no reduction in liver volume in *2-CLiP* larvae exposed to 0.10-2.00 µM Selinexor from 5 to 7 dpf (Fig. 8c,d) (Supplementary Fig. 6), *ahctf1^+/+^;TO(kras^G12V^)^T/+^* larvae exhibited a dose-dependent reduction in liver volume when exposed to Selinexor. Strikingly, liver volume was reduced even further in the context of *ahctf1* heterozygosity, such that at a 1.00 µM dose of Selinexor, *kras^G12V^*-driven hepatocyte hyperplasia was completely blocked and liver volume was indistinguishable from that of normal, non-hyperplastic *2-CLiP* larvae (Fig. 8c,d). These results indicate that XPO1 inhibition is an effective suppressor of growth and proliferation in our model of mutant Kras-driven HCC, and that its impact can be reinforced by a concomitant reduction in Elys function.

We next examined whether there was any clinical rationale for developing Nup inhibitors for the purpose of cancer treatment. We analysed the gene expression data of 372 HCC patients in the TCGA liver hepatocellular carcinoma (LIHC) dataset collated in the cBioPortal for Cancer Genomics^39,40^ for *AHCTF1* and other genes encoding components of the NUP107-160 complex (Supplementary Table 3). In a striking result, we found that the median overall survival of patients with high mRNA expression of components of the NUP107-160 complex, including ELYS is 21.70 months, whereas the survival of patients with low expression of mRNAs encoding NUP107-160 components is markedly extended to 81.73 months (Fig. 8e). Since the 372 samples in the LIHC dataset were profiled across 19 sequencing plates that could cause batch effects and unwanted variation, we determined the integrity of this result by performing ANOVA tests to assess the effects of individual sequencing plates on the expression of genes encoding components of the NUP107-160 complex (Supplementary Fig. 7). These analyses revealed that differences in gene expression observed between samples were largely unaffected by unwanted variation between sequencing plates. This clinical observation supports our proposition that ELYS and potentially other Nups would be useful targets for cancer treatment.

## DISCUSSION

In this paper we show that *ahctf1* heterozygosity markedly reduces tumour burden in a genetically engineered zebrafish model of HCC. Several factors indicate that this model provides a clinically relevant platform for the study of human HCC and the discovery of new therapeutic strategies^24^. Firstly, RAS/RAF/MAPK signalling is almost always hyper-activated in human HCC^41^. Secondly, the progressive accumulation of histopathologic features including irregular nuclei, cytoplasmic vacuolation and increased vascularisation that are characteristic of tumour progression in HCC are reproduced in the zebrafish model^42^. Thirdly, comparative transcriptomic analysis reveals that the gene expression profile exhibited by oncogenic *kras^G12V^* hepatocytes isolated from larval zebrafish strongly resembles that of early-stage human HCC, with elevated expression of RAS/RAF/MAPK target genes such as *FGFR4, ETV4, EPHA2, DUSP6* and *SPRY* and DNA damage response genes *GADD45B, CCND1* and *H2AX1*^43^.

Our results with this *in vivo* model of HCC compliment a growing body of evidence that carcinogenesis places persistently high demands on essential cellular genes, including those required for proper nuclear pore function, creating a vulnerability that may be therapeutically targeted without causing adverse effects on normal tissue^44,45^. For example, Sakuma *et al*. used siRNAs targeting 28 out of 32 Nup genes, to reduce the abundance of mature NPCs in the human melanoma-derived cell line, A375^44^. Focusing on two Nups exhibiting a severe reduction in NPC density, this study showed that NUP160 and NUP93 are essential for NPC assembly and that inhibiting this process results in cell death. Moreover, they found that cancer cells exhibited heightened susceptibility to inhibition of NPC assembly compared to human pulmonary fibroblasts and a normal cell line derived from retinal pigmental epithelium (RPE1), which instead underwent reversible cell cycle arrest. The differential sensitivity between cancer cells and normal cells to NPC assembly inhibition is consistent with reports that quiescent cells in normal tissues generally maintain low levels of Nup expression and may exist for many years with the same set of assembled NPCs^46,47^.

That *AHCTF1* might play a role in *RAS*-fuelled cancer cells was first indicated in a genome-wide RNAi screen carried-out in human colorectal cancer-derived DLD1 cells^3^. Since then, a collection of genome-wide CRISPR knockout screens has shown that *AHCTF1* is essential for the survival of 786/789 human cancer cell lines^48,49^ (Supplementary Table 3). While these data show that *AHCTF1* is indispensable for virtually all cancer cell lines, *AHCTF1* expression is severely and irreversibly depleted in this *in vitro* setting (>90%), and there is no read-out of how this degree of knockdown impacts on healthy cells. We believe our *in vivo* experiments are more informative, not only because they show that a 50% reduction in *ahctf1* gene dosage markedly reduces tumour burden but also because they show that this is achieved without a detrimental impact on healthy cells.

Mutations in *TP53*, or amplification/overexpression of its negative regulators *MDM2*/*MDM4*, occur in 30% of HCC cases^50^. This is very relevant to our study because we show that a prominent response to *ahctf1* heterozygosity in our *in vivo* HCC model is accumulation of DNA damage and activation of Tp53. We therefore wondered whether the potential therapeutic effect of inhibiting ELYS function in a clinical cancer setting would be diminished in tumours lacking functional TP53 protein. To address this question, we studied the impact of *ahctf1* heterozygosity in our *in vivo* HCC model on a homozygous *tp53* mutant background. We found that *ahctf1* heterozygosity still produced a 24% reduction in tumour burden and induced cell death in the complete absence of Tp53 function, leading us to consider whether the loss of Tp53 function was compensated for by increased *tp63* and/or *tp73* expression. These genes encode Tp53 family proteins that share considerable structural homology with Tp53, particularly within the DNA binding domain, and they can activate common and distinct sets of target genes to produce cell cycle arrest, senescence and apoptosis^51,52^. For example, in response to DNA damage, Tp63 induces senescence and apoptosis in the same way as Tp53, via transcriptional induction of *cdkn1a*, *bbc3* and *pmaip1*^53^. We found that *ahctf1* heterozygosity upregulated *tp63* and *tp73* expression by 3.0-fold and 2.3-fold, respectively, and that this was further enhanced by homozygous *tp53* mutation. These results suggest a mechanism through which *ahctf1* heterozygosity achieves a reduction in tumour burden in the absence of Tp53 function. Since *TP63* and *TP73* are rarely mutated in cancer^54^, we think that inhibition of ELYS function is likely to produce a beneficial effect on tumour growth, even in patients harbouring *TP53* mutations.

Dysregulation of nucleocytoplasmic transport is a common feature in a broad spectrum of cancers, usually arising from altered expression of nuclear pore components and/or nuclear transport receptors^55^. For example, overexpression of POM121, a transmembrane NPC component, enhances the nuclear import of the pro-proliferation transcription factors, E2F1, MYC and AR, and increases the therapeutic resistance of prostate cancer^56^. Similarly, the nuclear transport receptor XPO1 is frequently overexpressed in cancers, including HCC, and in this case leads to mis-localisation and inactivation of tumour suppressor proteins^38,57^. As a result, numerous selective inhibitors of nuclear export (SINEs) have been developed, including Selinexor (KPT-330), which has advanced through clinical trials and received FDA approval for the treatment of relapsed or refractory multiple myeloma and diffuse large B-cell lymphoma^37^. We found that Selinexor treatment in our zebrafish HCC model produced a dose-dependent reduction in tumour burden. Moreover, this therapeutic effect was strongly augmented by *ahctf1* heterozygosity, suggesting that new drugs targeting the ELYS protein could be combined effectively with drugs that target nucleocytoplasmic transport. We also exploited the genetic tractability of our zebrafish HCC model to show that trans heterozygosity of *ahctf1*, *ranbp2* delivers a synergistic reduction in liver volume/tumour burden, bringing it back to the volume of non-hyperplastic livers. Notably, non-*TO(kras^G12V^)* livers were completely unaffected by *ahctf1*, *ranbp2* trans heterozygosity, indicating a three-way synthetic lethal interaction in cancer cells that may be possible to recapitulate with combinatorial drug treatments.

The dynamic and diverse functions of ELYS require its binding to Nups and many other proteins, including components of kinetochore and replicative licensing complexes, as well as binding to chromatin in various states of condensation. The molecular topology of the ELYS protein and the structural basis for its diverse protein-protein and protein-chromatin interactions are hitherto poorly defined. Crystal structures of ELYS have shown that the *β*-propeller and *α*-helical domains in the N-terminal half of the ELYS protein bind to NUP160, and cryo-electron microscopy reveals the association of the ELYS C-terminal peptide with nucleosomes^21,58^. However, further high-resolution structural characterization is required to advance our molecular understanding of the extent to which discrete domains within the ELYS protein contribute to its various interactions, and to reveal opportunities to inhibit such regions for the purpose of disabling ELYS function therapeutically. It is possible that the topology of ELYS lacks features that would facilitate high affinity binding with small drug-like molecules, as has been suggested for other Nups with scaffolding properties^44^. However, rapidly emerging technological advances in drug development are likely to provide alternative strategies to target the ELYS protein. For example, carefully controlled proteasomal degradation of proteins that participate in the oncogenic process using proteolysis targeting chimeras (PROTACs) is a rapidly advancing field in cancer therapy^59^. This approach depends on the availability of a ligand/probe to direct the PROTAC to the targeted protein, but this need not be a druggable site of protein-protein or protein-chromatin interaction, as is the case for small molecules designed to disrupt the function of the protein. In parallel, emerging small molecule RNA-targeting technology is on course to permit direct modulation of the abundance of specific RNA transcripts for a variety of clinical purposes^60^. In summary, we believe our *in vivo* cancer studies provide a strong and feasible rationale for targeting ELYS for the treatment of a broad spectrum of cancers and suggest potential avenues for effective combinatorial treatments.

## MATERIALS AND METHODS

### Zebrafish maintenance and strains

Zebrafish were maintained at 28°C on a 12 h light/12 h dark cycle. The mutant lines *ahctf1^ti262^* and *tp53^M214K/M214K^* (also known as *tp53^e7/e7^* and referred to herein as *tp53^m/m^*) have been described previously^7,26^. Tg(*fabp10:dsRed, ela3l:GFP*)^gz12^, hereafter referred to as *2-CLiP*, expresses dsRed in the liver and GFP in the exocrine pancreas but carries no oncogenic transgenes or mutations^25^. The Tg(*fabp10:rtTA2s-M2;TRE2:EGFP-kras^G12V^*)^gz32^ line referred to as *TO(kras^G12V^)^T/+^* and the cell death reporter line, Tg(*actb2:SEC-Hsa.ANXA5-mKate2,cryaa:mCherry*)^uq24rp^ were described previously^24,27^. The *ranbp2^s452^* mutant line was generated in the Liver^plus^ ENU mutagenesis screen^32^, and its genetic and morphological characterization is presented in this paper.

### Inducing hepatocyte hyperplasia in transgenic zebrafish

To induce *kras^G12V^* expression, *TO(kras^G12V^)^T/+^* embryos were treated with 20 µg/mL doxycycline (Sigma, #D9891) at 2 dpf in E3 medium (5 mM NaCl, 0.17 mM KCl, 0.33 mM CaCl2, 0.33 mM MgSO4) with 0.003% 1-Phenyl-2-thiourea (PTU; Sigma, #P7629) to suppress pigmentation. E3 medium was changed at 5 dpf and fresh doxycycline (final conc. 20 µg/mL) added. For the Selinexor experiments, the drug (final concentration 0.10-2.00 μM) (Karyopharm) or 0.001% DMSO (vehicle control) was added at 5 dpf. Morphological and molecular analyses were performed at 7 dpf. To quantitate liver volume, larvae were anaesthetized with benzocaine (200 mg/L) and mounted in 1% agarose. Image acquisition was performed using an Olympus FVMPE-RS multiphoton microscope with a 25x objective and Olympus FV30-SW software. Excitation wavelengths for GFP and RFP were 840 nm and 1100 nm, respectively. For volumetric analysis of whole livers, z-stacks with step-size 2 μm, were imported into ImageJ or Imaris software.

### Genotyping

Genomic DNA (gDNA) was extracted from whole zebrafish larvae by incubation at 95°C in 50 μL of 50 mM sodium hydroxide (NaOH) for 20 min, followed by neutralization with 5 μL of 1 M Tris-HCl (pH 8.0). Primer sequences for genotyping are listed in Supplementary Table 1.

### mRNA expression analysis

Total RNA was extracted from independent pools of micro-dissected zebrafish livers using the RNeasy Micro Kit (QIAGEN, #74004). RNA integrity was assessed by a High Sensitivity RNA ScreenTape assay (Agilent, #5067-5579) on a 2200 TapeStation. cDNA was generated from 1-10 µg RNA using the Superscript III First Strand Synthesis System (Invitrogen, #18080051) and oligo(dT) priming according to the manufacturer’s instructions. RT-quantitative PCR (RT-qPCR) was performed using a SensiMix SYBR kit (Bioline, #QT605-05) on an Applied Biosystems ViiATM7 Real-Time PCR machine. Expression data were normalized by reference to *hrpt1*, *b2m* and *tbp*. LinRegPCR V11.0 was used for baseline correction, PCR efficiency calculation and transcript quantification analysis^61^. Relative expression levels were calculated by the 2^−ΔΔCt^ method and all results were expressed as the mean ± SEM of three independent pools of biological replicates. Primer sequences are listed in Supplementary Table 2.

### Western blot analysis

Pooled micro-dissected zebrafish livers were lysed in RIPA buffer (20mM Hepes, pH 7.9, 150mM NaCl, 1mM MgCl2, 1% NP40, 10mM NaF, 0.2mM Na3VO4, 10mM *β*-glycerol phosphate) supplemented with cOmplete Proteinase Inhibitor (Roche, #11836170001) and PhosTOP phosphatase inhibitors (Roche, #04906837001). Samples were incubated for 30 min on ice and the extracts cleared by centrifugation at 13,000 rpm for 20 min at 4°C. The protein concentration of samples was determined by BCA protein assay (Thermo Fisher Scientific, #23227). 25 μg of protein per lane were resolved on NuPAGE Novex Bis-Tris 4-12% polyacrylamide gels (Invitrogen, #NP0321BOX) and transferred onto nitrocellulose blotting membranes (Amersham Protran, #10600003). Membranes were blocked with 5% BSA in PBS for 1 h at RT and incubated with primary antibodies 1:500 Anti-p53 (9.1) mouse mAb (Abcam, #ab77813) overnight at 4°C and 1:1000 Anti-GAPDH (14C10) Rabbit mAb (Cell Signalling Technology, #2118) for 1 h at RT. Secondary antibodies: Goat anti-mouse HRP (Dako, #P0447) and Goat anti-Rabbit HRP (Dako, #P0448), were used at 1:5000 and incubated with membranes for 1 h at RT. Signals were developed using Amersham ECL Western Blotting Detection Kit (Cytiva, # RPN2108) and imaged on a Chemidoc Touch (Biorad). Relative protein quantitation was calculated based on normalized integrated intensity.

### Cell death analysis

To assess apoptosis, 7 dpf *TO(kras^G12V^)*;*Annexin 5-mKate* zebrafish larvae were fixed in 4% paraformaldehyde (PFA) overnight at 4°C and livers isolated by micro-dissection. Image acquisition was performed using a Zeiss LSM 880 microscope with a 20x objective and ZEN software. Excitation wavelengths for mKate and GFP were 560 nm and 900 nm, respectively. Liver volume was quantified and 3D segmentation of Annexin 5-mKate signals were performed in ImageJ.

### Cell cycle analysis

EdU incorporation was used to assess the percentage of cells in S phase of the cell cycle. Briefly, live zebrafish larvae (7 dpf) were incubated at 28°C in 2 mM EdU (Invitrogen, #C10340) in E3 medium for 2 h followed by a further incubation in fresh E3 medium for 1 h. Larvae were euthanised using benzocaine (1000 mg/L; Sigma, #PHR1158) prior to removal of the liver by micro-dissection. EdU labelling was carried out using the Click-iT Edu Alexa Fluor 647 (AF647) imaging kit (Invitrogen, #C10340), according to the manufacturer’s instructions. The livers were co-stained with Hoechst 33342 (Thermofisher, #62249). Image acquisition was performed using an Olympus FVMPE-RS multiphoton microscope with excitation wavelengths of 950 nm for Hoechst 33342 dye and 1160 nm for AF647 (Thermofisher, #A21235). The number of Hoechst 33342 and EdU positive cells was quantified using Arivis Vision4D software.

### Nuclear pore analysis

Larvae fixed in 4% PFA overnight at 4°C were embedded in 4% low melting temperature agarose and transverse sections collected at 200 µm intervals using a vibrating microtome (Leica VT 1000S). Sections were blocked with 1% BSA in PBS/0.3% Triton X-100 and incubated with a 1:750 dilution of mAb414 (Abcam, #ab24609) at 4°C overnight. Sections were then incubated with 1:500 anti-mouse AF647 and Hoechst 33342 at room temperature for 1 h. Sections were mounted and imaged using a Zeiss LSM880 Fast Airyscan Confocal microscope with a 63x objective. 3D segmentation of cells was performed in ImageJ using EGFP signal for *TO(kras^G12V^)^T/+^* larvae or F-actin stained with 1:200 rhodamine phalloidin for *TO(kras^G12V^)^+/+^* larvae. An outline was drawn around the nuclear periphery and segmented using the Hoechst signal. mAb414 fluorescence intensity was calculated for the nuclear periphery and for the cytoplasm. NPC density was calculated by finding maxima for mAb414 at the nuclear surface of 5 nuclei per liver.

### Cryosectioning and immunofluorescence microscopy analysis

Dissected livers were fixed in 4% PFA overnight at 4°C and washed with PBS/0.1% Tween 20 before incubation in 30% sucrose in PBS overnight at 4°C. Livers were aligned in a tissue mould, embedded in OCT and frozen on dry ice. The livers were sectioned at 10 µm intervals using a Thermofisher Scientific Microm HM550 cryostat. Sections were washed with PBS before blocking with 10% FCS in PBS/0.3% Triton X-100. Incubation with primary antibodies was performed at 4°C overnight, while incubation with secondary antibodies was performed at room temperature for 1 h. Antibodies used in this work were: 1:2000 *α*-Tubulin DM1A (CST, #3873), 1:1000 γ-H2AX (gift of James Amatruda), 1:250 cleaved caspase 3 (CST, #9664), 1:500 anti-rabbit AF647 (Thermofisher scientific, #A31573) and 1:500 anti-mouse AF647 (Thermofisher, #A21235). Prolong Diamond Antifade reagent with DAPI (Thermofisher #P36962) was used for slide mounting. A Zeiss LSM880 Fast Airyscan Confocal microscope with a 63x objective was used for image acquisition and ImageJ for image analysis.

### Statistical analysis

Data are expressed as mean ± SEM unless indicated otherwise and the number of biological replicates indicating samples from individual animals/livers, or pools of individual animals/livers for each experiment are stated in the figure legends. P-values were calculated using Student’s *t*-tests (two-tailed, followed by Welch’s correction) when comparing two groups, and by ANOVA followed by Tukey’s post-hoc test when comparing multiple groups. The effect of Selinexor treatment on liver volume was analysed by linear regression, regressing liver volume against Selinexor concentration. All analysis was performed using GraphPad Prism V7.03 (GraphPad software) and p-values ≤ 0.05 were considered statistically significant.

## ACKNOWLEDGEMENTS

The authors thank Tyson Blanch, Cameron Mackey, Elizabeth Grgacic and Bryan Ko (zebrafish husbandry), Ellen Tsui (histology), James Amatruda (rabbit polyclonal antibody to zebrafish *γ*-H2AX), Brendon Monahan and Leigh Coultas (insightful discussions).

## ADDITIONAL INFORMATION

### AUTHORS’ CONTRIBUTIONS

KM, KD, JKH conceived and designed the experiments; KM, KD, KS, BH, CS, GB, RM, ATP, TH, EO, DYRS, ZG, JKH developed the methodology; KM, KD, FG, LW, KS, BH, CS, GB, RM, ATP, JKH acquired and analysed the data; KM, KD and JKH wrote the manuscript; KD, JKH supervised the study; JKH acquired the funding.

### DATA AVAILABILITY

The datasets generated and/or analysed during the current study are available in the cBioPortal Cancer Genomics database (http://www.cbioportal.org).

## REFERENCES

1 Gao, S. & Lai, L. Synthetic lethality in drug development: the dawn is coming. Future Med Chem 10, 2129–2132, doi:10.4155/fmc-2018-0227 (2018).

2 Lord, C. J. & Ashworth, A. PARP inhibitors: Synthetic lethality in the clinic. Science 355, 1152–1158, doi:10.1126/science.aam7344 (2017).

3 Luo, J. et al. A genome-wide RNAi screen identifies multiple synthetic lethal interactions with the Ras oncogene. Cell 137, 835–848, doi:10.1016/j.cell.2009.05.006 (2009).

4 Wang, T. et al. Gene Essentiality Profiling Reveals Gene Networks and Synthetic Lethal Interactions with Oncogenic Ras. Cell 168, 890–903 e815, doi:10.1016/j.cell.2017.01.013 (2017).

5 Solimini, N. L., Luo, J. & Elledge, S. J. Non-oncogene addiction and the stress phenotype of cancer cells. Cell 130, 986–988, doi:10.1016/j.cell.2007.09.007 (2007).

6 Okita, K. et al. Targeted disruption of the mouse ELYS gene results in embryonic death at peri-implantation development. Genes Cells 9, 1083–1091, doi:10.1111/j.1365-2443.2004.00791.x (2004).

7 Chen, J. N. et al. Mutations affecting the cardiovascular system and other internal organs in zebrafish. Development 123, 293–302 (1996).

8 de Jong-Curtain, T. A. et al. Abnormal nuclear pore formation triggers apoptosis in the intestinal epithelium of elys-deficient zebrafish. Gastroenterology 136, 902–911, doi:10.1053/j.gastro.2008.11.012 (2009).

9 Davuluri, G. et al. Mutation of the zebrafish nucleoporin elys sensitizes tissue progenitors to replication stress. PLoS Genet 4, e1000240, doi:10.1371/journal.pgen.1000240 (2008).

10 Rasala, B. A., Orjalo, A. V., Shen, Z., Briggs, S. & Forbes, D. J. ELYS is a dual nucleoporin/kinetochore protein required for nuclear pore assembly and proper cell division. Proc Natl Acad Sci U S A 103, 17801–17806, doi:10.1073/pnas.0608484103 (2006).

11 Gillespie, P. J., Khoudoli, G. A., Stewart, G., Swedlow, J. R. & Blow, J. J. ELYS/MEL-28 chromatin association coordinates nuclear pore complex assembly and replication licensing. Curr Biol 17, 1657–1662, doi:10.1016/j.cub.2007.08.041 (2007).

12 Franz, C. et al. MEL-28/ELYS is required for the recruitment of nucleoporins to chromatin and postmitotic nuclear pore complex assembly. EMBO Rep 8, 165–172, doi:10.1038/sj.embor.7400889 (2007).

13 Beck, M. & Hurt, E. The nuclear pore complex: understanding its function through structural insight. Nat Rev Mol Cell Biol 18, 73–89, doi:10.1038/nrm.2016.147 (2017).

14 Hodgson, B., Li, A., Tada, S. & Blow, J. J. Geminin becomes activated as an inhibitor of Cdt1/RLF-B following nuclear import. Curr Biol 12, 678–683, doi:10.1016/s0960-9822(02)00778-9 (2002).

15 Kuhn, T. M., Pascual-Garcia, P., Gozalo, A., Little, S. C. & Capelson, M. Chromatin targeting of nuclear pore proteins induces chromatin decondensation. J Cell Biol 218, 2945–2961, doi:10.1083/jcb.201807139 (2019).

16 Guttinger, S., Laurell, E. & Kutay, U. Orchestrating nuclear envelope disassembly and reassembly during mitosis. Nat Rev Mol Cell Biol 10, 178–191, doi:10.1038/nrm2641 (2009).

17 Chatel, G. & Fahrenkrog, B. Nucleoporins: leaving the nuclear pore complex for a successful mitosis. Cell Signal 23, 1555–1562, doi:10.1016/j.cellsig.2011.05.023 (2011).

18 Mishra, R. K., Chakraborty, P., Arnaoutov, A., Fontoura, B. M. & Dasso, M. The Nup107-160 complex and gamma-TuRC regulate microtubule polymerization at kinetochores. Nat Cell Biol 12, 164–169, doi:10.1038/ncb2016 (2010).

19 Orjalo, A. V. et al. The Nup107-160 nucleoporin complex is required for correct bipolar spindle assembly. Mol Biol Cell 17, 3806–3818, doi:10.1091/mbc.e05-11-1061 (2006).

20 Yokoyama, H. et al. The nucleoporin MEL-28 promotes RanGTP-dependent gamma-tubulin recruitment and microtubule nucleation in mitotic spindle formation. Nat Commun 5, 3270, doi:10.1038/ncomms4270 (2014).

21 Kobayashi, W. et al. Structural and biochemical analyses of the nuclear pore complex component ELYS identify residues responsible for nucleosome binding. Commun Biol 2, 163, doi:10.1038/s42003-019-0385-7 (2019).

22 Bilokapic, S. & Schwartz, T. U. Molecular basis for Nup37 and ELY5/ELYS recruitment to the nuclear pore complex. Proc Natl Acad Sci U S A 109, 15241–15246, doi:10.1073/pnas.1205151109 (2012).

23 Rasala, B. A., Ramos, C., Harel, A. & Forbes, D. J. Capture of AT-rich chromatin by ELYS recruits POM121 and NDC1 to initiate nuclear pore assembly. Mol Biol Cell 19, 3982–3996, doi:10.1091/mbc.E08-01-0012 (2008).

24 Chew, T. W. et al. Crosstalk of Ras and Rho: activation of RhoA abates Kras-induced liver tumorigenesis in transgenic zebrafish models. Oncogene 33, 2717–2727, doi:10.1038/onc.2013.240 (2014).

25 Korzh, S. et al. Requirement of vasculogenesis and blood circulation in late stages of liver growth in zebrafish. BMC Dev Biol 8, 84, doi:10.1186/1471-213X-8-84 (2008).

26 Berghmans, S. et al. tp53 mutant zebrafish develop malignant peripheral nerve sheath tumors. Proc Natl Acad Sci U S A 102, 407–412, doi:10.1073/pnas.0406252102 (2005).

27 Hall, T. E. et al. Cellular rescue in a zebrafish model of congenital muscular dystrophy type 1A. NPJ Regen Med 4, 21, doi:10.1038/s41536-019-0084-5 (2019).

28 Joseph, J., Liu, S. T., Jablonski, S. A., Yen, T. J. & Dasso, M. The RanGAP1-RanBP2 complex is essential for microtubule-kinetochore interactions in vivo. Curr Biol 14, 611–617, doi:10.1016/j.cub.2004.03.031 (2004).

29 Salina, D., Enarson, P., Rattner, J. B. & Burke, B. Nup358 integrates nuclear envelope breakdown with kinetochore assembly. J Cell Biol 162, 991–1001, doi:10.1083/jcb.200304080 (2003).

30 Hashizume, C., Kobayashi, A. & Wong, R. W. Down-modulation of nucleoporin RanBP2/Nup358 impaired chromosomal alignment and induced mitotic catastrophe. Cell Death Dis 4, e854, doi:10.1038/cddis.2013.370 (2013).

31 Vecchione, L. et al. A Vulnerability of a Subset of Colon Cancers with Potential Clinical Utility. Cell 165, 317–330, doi:10.1016/j.cell.2016.02.059 (2016).

32 Ober, E. A., Verkade, H., Field, H. A. & Stainier, D. Y. Mesodermal Wnt2b signalling positively regulates liver specification. Nature 442, 688–691, doi:10.1038/nature04888 (2006).

33 Leshchiner, I. et al. Mutation mapping and identification by whole-genome sequencing. Genome Res 22, 1541–1548, doi:10.1101/gr.135541.111 (2012).

34 Voz, M. L. et al. Fast homozygosity mapping and identification of a zebrafish ENU-induced mutation by whole-genome sequencing. PLoS One 7, e34671, doi:10.1371/journal.pone.0034671 (2012).

35 Boglev, Y. et al. Autophagy induction is a Tor-and Tp53-independent cell survival response in a zebrafish model of disrupted ribosome biogenesis. PLoS Genet 9, e1003279, doi:10.1371/journal.pgen.1003279 (2013).

36 Markmiller, S. et al. Minor class splicing shapes the zebrafish transcriptome during development. Proc Natl Acad Sci U S A 111, 3062–3067, doi:10.1073/pnas.1305536111 (2014).

37 Jans, D. A., Martin, A. J. & Wagstaff, K. M. Inhibitors of nuclear transport. Curr Opin Cell Biol 58, 50–60, doi:10.1016/j.ceb.2019.01.001 (2019).

38 Zheng, Y. et al. KPT-330 inhibitor of XPO1-mediated nuclear export has anti-proliferative activity in hepatocellular carcinoma. Cancer Chemother Pharmacol 74, 487–495, doi:10.1007/s00280-014-2495-8 (2014).

39 Cerami, E. et al. The cBio cancer genomics portal: an open platform for exploring multidimensional cancer genomics data. Cancer Discov 2, 401–404, doi:10.1158/2159-8290.CD-12-0095 (2012).

40 Gao, J. et al. Integrative analysis of complex cancer genomics and clinical profiles using the cBioPortal. Sci Signal 6, pl1, doi:10.1126/scisignal.2004088 (2013).

41 Calvisi, D. F. et al. Ubiquitous activation of Ras and Jak/Stat pathways in human HCC. Gastroenterology 130, 1117–1128, doi:10.1053/j.gastro.2006.01.006 (2006).

42 Zheng, W. et al. Xmrk, kras and myc transgenic zebrafish liver cancer models share molecular signatures with subsets of human hepatocellular carcinoma. PLoS One 9, e91179, doi:10.1371/journal.pone.0091179 (2014).

43 Huo, X. et al. Transcriptomic analyses of oncogenic hepatocytes reveal common and different molecular pathways of hepatocarcinogenesis in different developmental stages and genders in kras(G12V) transgenic zebrafish. Biochem Biophys Res Commun 510, 558–564, doi:10.1016/j.bbrc.2019.02.008 (2019).

44 Sakuma, S. et al. Inhibition of Nuclear Pore Complex Formation Selectively Induces Cancer Cell Death. Cancer Discov 11, 176–193, doi:10.1158/2159-8290.CD-20-0581 (2021).

45 Beck, M., Schirmacher, P. & Singer, S. Alterations of the nuclear transport system in hepatocellular carcinoma - New basis for therapeutic strategies. J Hepatol 67, 1051–1061, doi:10.1016/j.jhep.2017.06.021 (2017).

46 D’Angelo, M. A., Raices, M., Panowski, S. H. & Hetzer, M. W. Age-dependent deterioration of nuclear pore complexes causes a loss of nuclear integrity in postmitotic cells. Cell 136, 284–295, doi:10.1016/j.cell.2008.11.037 (2009).

47 Toyama, B. H. et al. Identification of long-lived proteins reveals exceptional stability of essential cellular structures. Cell 154, 971–982, doi:10.1016/j.cell.2013.07.037 (2013).

48 Dempster, J. M. et al. Extracting Biological Insights from the Project Achilles Genome-Scale CRISPR Screens in Cancer Cell Lines. doi:10.1101/720243 (2019).

49 Meyers, R. M. et al. Computational correction of copy number effect improves specificity of CRISPR-Cas9 essentiality screens in cancer cells. Nat Genet 49, 1779–1784, doi:10.1038/ng.3984 (2017).

50 Zucman-Rossi, J., Villanueva, A., Nault, J. C. & Llovet, J. M. Genetic Landscape and Biomarkers of Hepatocellular Carcinoma. Gastroenterology 149, 1226–1239 e1224, doi:10.1053/j.gastro.2015.05.061 (2015).

51 Levrero, M. et al. The p53/p63/p73 family of transcription factors: overlapping and distinct functions. J Cell Sci 113 (Pt 10), 1661–1670 (2000).

52 Suh, E. K. et al. p63 protects the female germ line during meiotic arrest. Nature 444, 624–628, doi:10.1038/nature05337 (2006).

53 Kerr, J. B. et al. DNA damage-induced primordial follicle oocyte apoptosis and loss of fertility require TAp63-mediated induction of Puma and Noxa. Mol Cell 48, 343–352, doi:10.1016/j.molcel.2012.08.017 (2012).

54 Dotsch, V., Bernassola, F., Coutandin, D., Candi, E. & Melino, G. p63 and p73, the ancestors of p53. Cold Spring Harb Perspect Biol 2, a004887, doi:10.1101/cshperspect.a004887 (2010).

55 Kau, T. R., Way, J. C. & Silver, P. A. Nuclear transport and cancer: from mechanism to intervention. Nat Rev Cancer 4, 106–117, doi:10.1038/nrc1274 (2004).

56 Rodriguez-Bravo, V. et al. Nuclear Pores Promote Lethal Prostate Cancer by Increasing POM121-Driven E2F1, MYC, and AR Nuclear Import. Cell 174, 1200–1215 e1220, doi:10.1016/j.cell.2018.07.015 (2018).

57 Hill, R., Cautain, B., de Pedro, N. & Link, W. Targeting nucleocytoplasmic transport in cancer therapy. Oncotarget 5, 11–28, doi:10.18632/oncotarget.1457 (2014).

58 Bilokapic, S. & Schwartz, T. U. Structural and functional studies of the 252 kDa nucleoporin ELYS reveal distinct roles for its three tethered domains. Structure 21, 572–580, doi:10.1016/j.str.2013.02.006 (2013).

59 Ocana, A. & Pandiella, A. Proteolysis targeting chimeras (PROTACs) in cancer therapy. Journal of experimental & clinical cancer research : CR 39, 189, doi:10.1186/s13046-020-01672-1 (2020).

60 Mukherjee, H. et al. PEARL-seq: A Photoaffinity Platform for the Analysis of Small Molecule-RNA Interactions. ACS Chem Biol 15, 2374–2381, doi:10.1021/acschembio.0c00357 (2020).

61 Ruijter, J. M. et al. Amplification efficiency: linking baseline and bias in the analysis of quantitative PCR data. Nucleic Acids Res 37, e45, doi:10.1093/nar/gkp045 (2009).

