## Supplementary Material for "Elys deficiency constrains Kras-driven tumour burden by amplifying oncogenic stress"

### Supplementary Data

**Comprises:**

**Supplementary Figure 1, related to Figure 2**

**Supplementary Figure 2, related to Figure 2**

**Supplementary Figure 3, related to Figure 2**

**Supplementary Figure 4, related to Figure 3**

**Supplementary Figure 5, related to Figure 8**

**Supplementary Figure 6, related to Figure 8**

**Supplementary Figure 7, related to Figure 8**

**Supplementary Table 1**

**Supplementary Table 2**

**Supplementary Table 3**

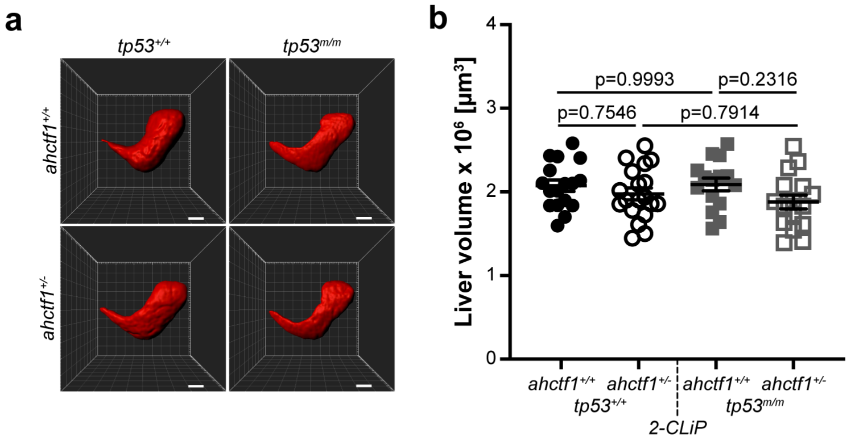


**Supplementary Fig. 1 *ahctf1* heterozygosity and/or homozygous *tp53* mutation do not impact on liver volume during normal liver development.**

**a.** Representative three-dimensional reconstructions of *2-CLiP* livers (not expressing a mutant *kras* transgene) of the indicated *ahctf1* and *tp53* genotypes. Scale bar 25 µm. **b.** Impact of *ahctf1* heterozygosity and *tp53* mutation on liver volume in *2-CLiP* larvae (n≥15). Data are expressed as mean ± SEM. Significance was assessed using a one-way ANOVA with Tukey's multiple comparisons test.

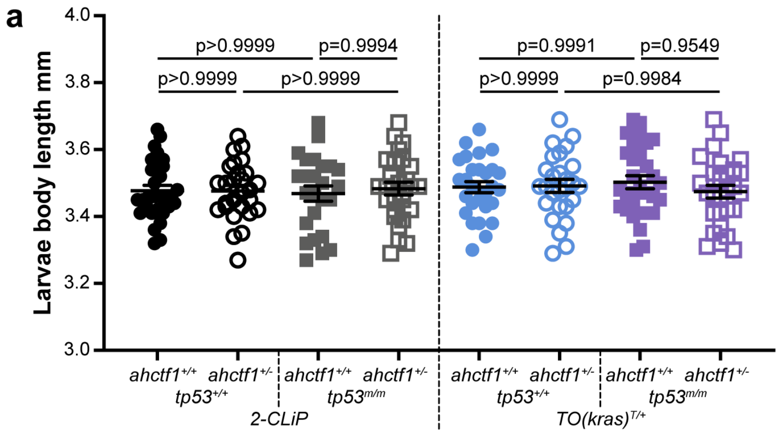


**Supplementary Fig. 2 *ahctf1* heterozygosity and/or homozygous *tp53* mutation do not impact on body length.**

**a.** Body length at 7 dpf in *2-CLiP* and *TO(kras$^{G12V}$)$^{T/+}$* larvae of the indicated *ahctf1* and *tp53* genotypes (n≥25). Data are expressed as mean ± SEM. Significance was calculated using a one-way ANOVA with Tukey's multiple comparisons test.

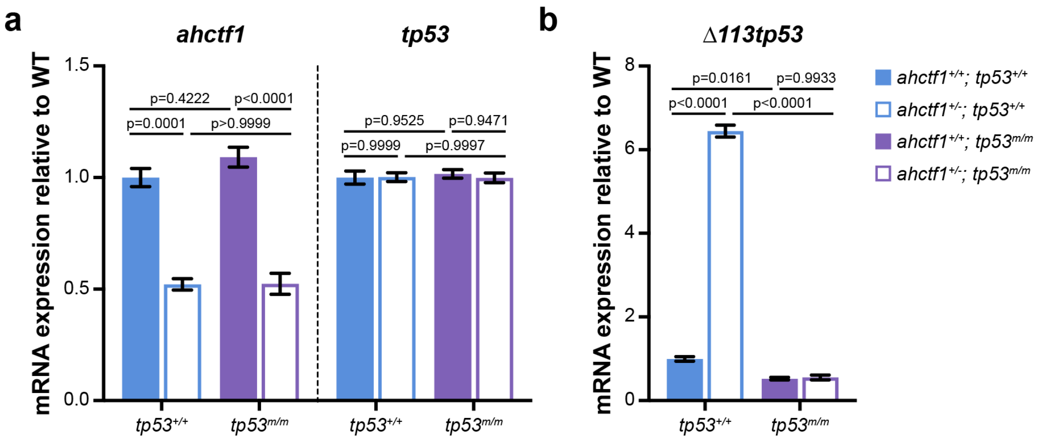


**Supplementary Fig. 3 *ahctf1* heterozygosity reduces *ahctf1* mRNA expression and increases expression of *Δ113tp53.***

**a.** RT-qPCR analysis of *ahctf1* and *tp53* mRNA expression and **b.** *Δ113tp53* (target of Tp53) mRNA expression in micro-dissected livers of *TO(kras$^{G12V}$)$^{T/+}$* larvae of the indicated *ahctf1* and *tp53* genotypes (n=3 biological replicates). Data are expressed as mean ± SEM. Significance was calculated using a one-way ANOVA with Tukey's multiple comparisons test.

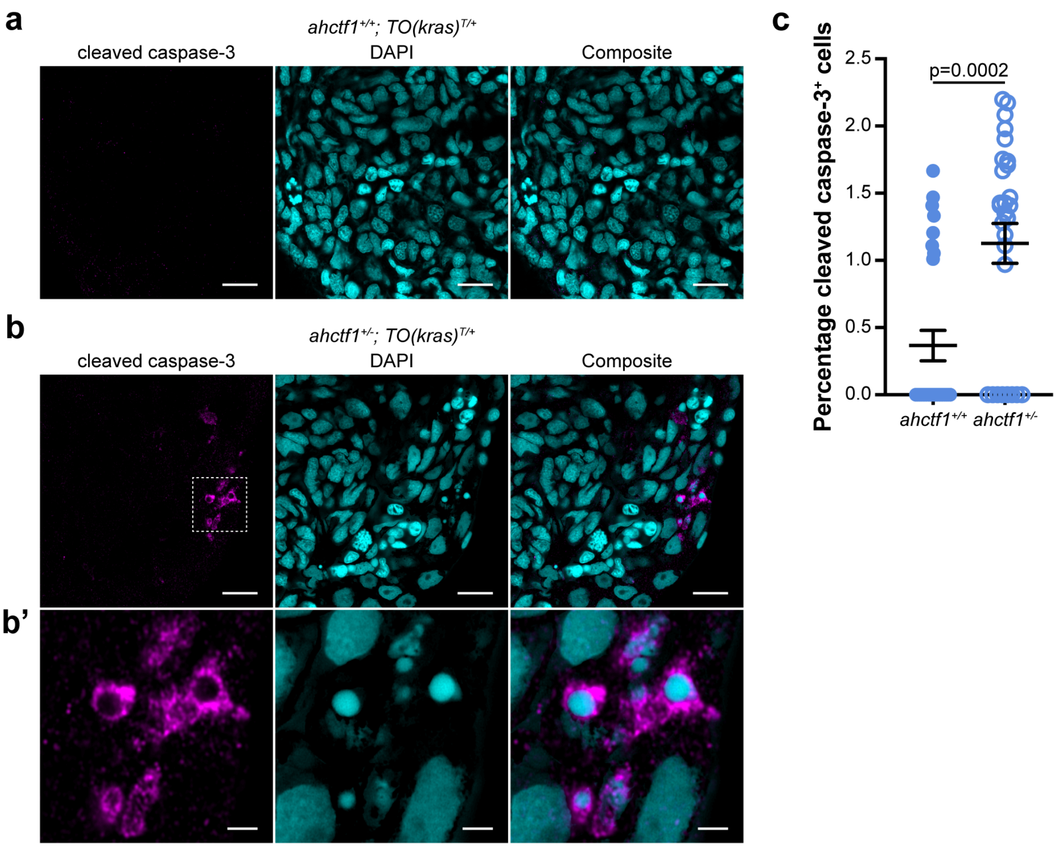


**Supplementary Fig. 4 *ahctf1* heterozygosity increases the number of *TO(kras$^{G12V}$)$^{T/+}$* hepatocytes expressing the cleaved, active form of caspase 3.**

**a.** Representative Airyscan images of *ahctf1$^{+/+}$;TO(kras$^{G12V}$)$^{T/+}$* liver cryosections stained with an antibody specific for the cleaved (active) form of caspase 3 (magenta) marking apoptosis and DAPI (cyan) marking DNA. Scale bar 5 µm **b.** Cryosections of liver from *ahctf1$^{+/-}$;TO(kras$^{G12V}$)$^{T/+}$* larvae reveal hepatocytes exhibiting cleaved caspase-3 staining and fragmented nuclei with condensed chromatin. Scale bar 5 µm. **b'.** Inset showing higher magnification image of cleaved caspase-3 positive hepatocytes in livers from *ahctf1$^{+/-}$;TO(kras$^{G12V}$)$^{T/+}$* larvae. Scale bar 2 µm. **c.** Quantification of the percentage of cells positive for active cleaved caspase-3 (n=28 liver sections). Data are expressed as mean ± SEM. Significance was calculated using a Student's t-test.

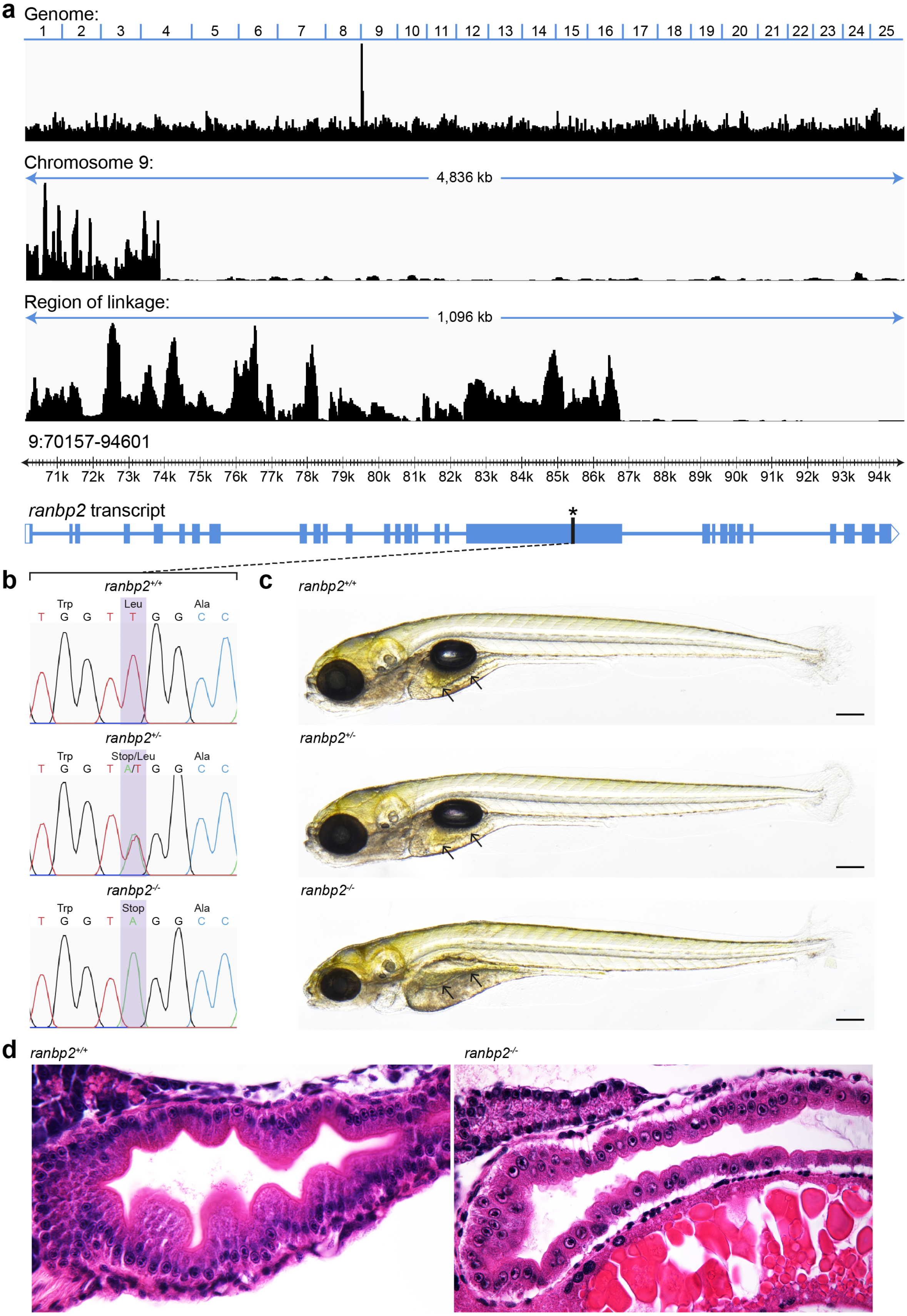


**Supplementary Fig. 5 Genetic and morphological characterisation of the *ranbp2* mutant.**

**a.** Positional cloning of the zebrafish mutant, *s452* identified in a transgene assisted ENU-mutagenesis screen for mutations affecting endodermal organ morphogenesis[32] using whole genome sequencing and homozygosity mapping. The genomic region containing the s452 causal mutation is derived exclusively from the initial ENU mutated allele and is defined by increased homozygosity. In *s452* mutants the region of linkage is restricted to a <1.1kb region on chromosome 9 within the *ranbp2* gene. Plot displays genomic homozygosity across 25 chromosomes, chromosome 9 and the region of linkage. The genome position of *ranbp2* within this region and a schematic diagram of the *ranbp2* transcript with an asterisk denoting the position of the mutation are also shown. **b.** Sanger sequencing chromatographs reveal a T>A transversion mutation introducing a premature stop codon (TTG>TAG). **c.** Brightfield images reveal abnormalities in the gross morphology of homozygous mutant *ranbp2* larvae at 7 dpf. *ranbp2*$^{-/-}$ larvae exhibit a thinner, flatter intestinal epithelium compared to wildtype and heterozygous *ranbp2* larvae (arrows). They also exhibit a smaller jaw, smaller eyes, failure to inflate the swim bladder and delayed resorption of the yolk. Scale bar 200 µm. **d.** Sagittal histological sections of 5 dpf *ranbp2* larvae stained with haematoxylin and eosin. Intestinal epithelial cells in *ranbp2*$^{-/-}$ larvae are smaller, unpolarised and the epithelium lacks characteristic folding, indicative of a proliferation defect.

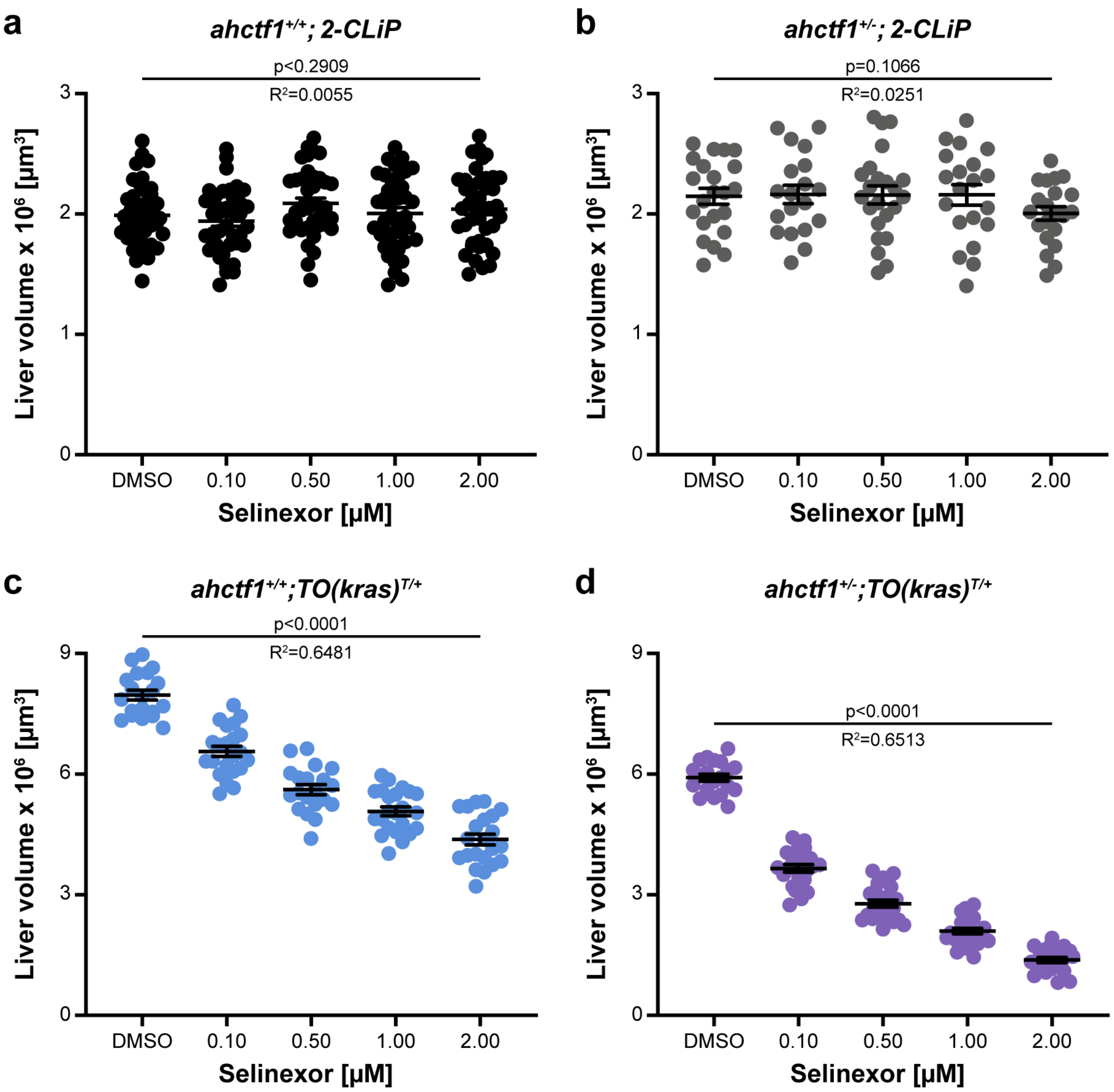


**Supplementary Fig. 6 Combination of Selinexor treatment and *ahctf1* heterozygosity reduces liver volume in *TO(kras$^{G12V}$)$^{T/+}$* larvae but has no impact on liver volume in *2-CLiP* larvae.**

Impact of Selinexor treatment (0.10-2.00 µM) on liver volume in **a.** *ahctf1$^{+/+}$;2-CLiP* larvae (n≥40). **b.** *ahctf1$^{+/-}$;2-CLiP* larvae (n≥19). **c.** *ahctf1$^{+/+}$;TO(kras$^{G12V}$)$^{T/+}$* larvae (n≥20). **d.** *ahctf1$^{+/-}$;TO(kras$^{G12V}$)$^{T/+}$* larvae (n>20). Data are expressed as mean ± SEM. Significance was calculated using a one-way ANOVA with Tukey's multiple comparisons test and linear regression analysis.

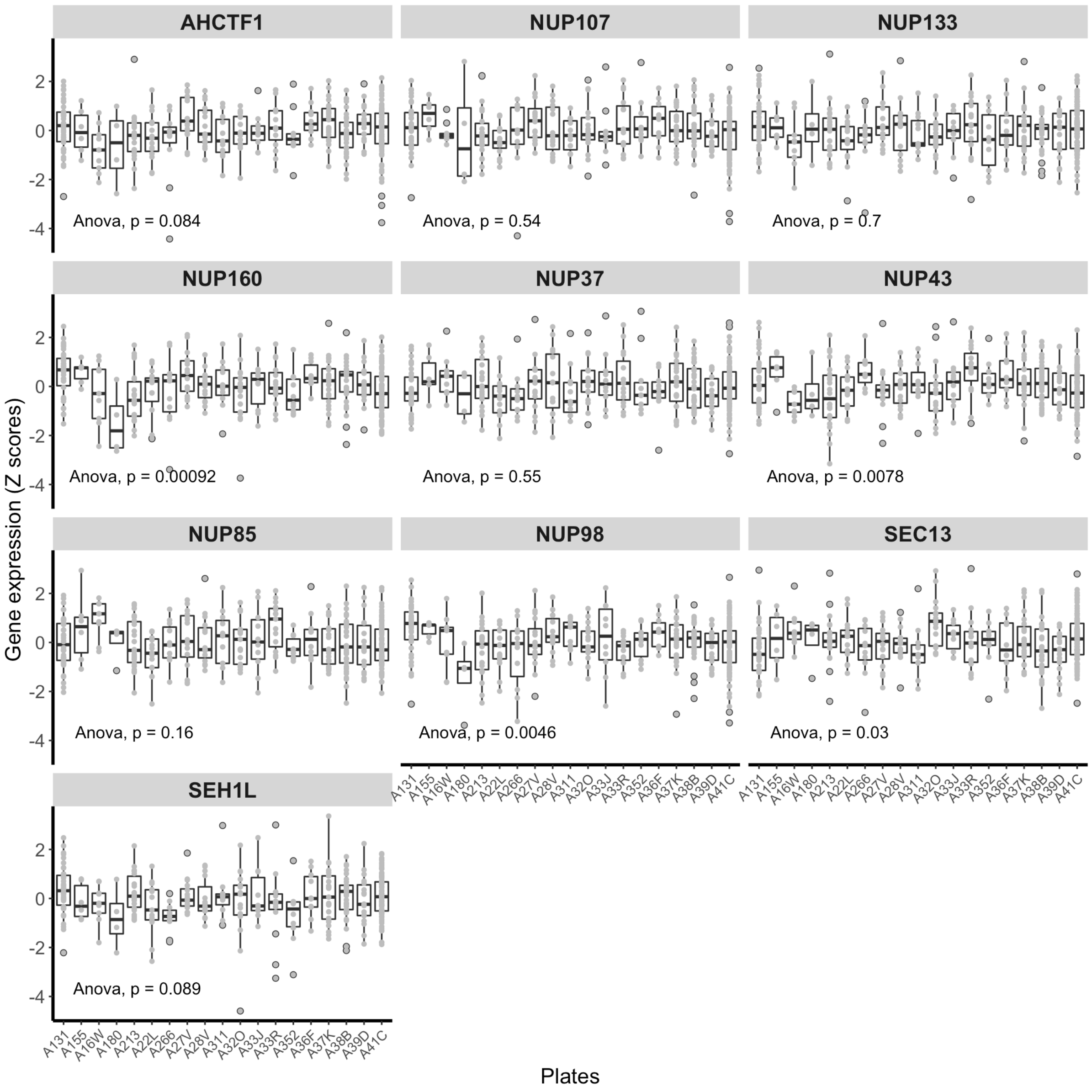


**Supplementary Fig. 7 NUP107-160 complex gene expression in the TCGA Liver hepatocellular carcinoma (LIHC) dataset is largely unaffected by unwanted variation.**

The expression patterns of genes in the NUP107-160 complex across sequencing plates in the TCGA LIHC RNA-Seq data. The X axes show the IDs for the 19 sequencing plates, and the Y axes represent Z scores of gene expression. The differences between average expression across individual sequencing plates were tested using ANOVA.

**Supplementary Table 1. Genotyping primers**

| Name | Forward (5'-3') | Reverse (5'-3') |
|---|---|---|
| *ahctf1*$^{ti262}$ | TGACATGCATGCCCTCTCTG | TAGCTGCTCCTCGCTTACGT |
| *ranbp2*$^{s452}$ | CGCCGATCAAGAGGACGAAA | TGTCCGCCGTAACACTACTC |
| *tp53* wildtype | AGCTGCATGGGGGGGAT | GATAGCCTAGTGCGAGCACACTCTT |
| *tp53* mutant | AGCTGCATGGGGGGGAA | GATAGCCTAGTGCGAGCACACTCTT |

**Supplementary Table 2. RT-qPCR primers**

| Name | Forward (5'-3') | Reverse (5'-3') |
|---|---|---|
| *ahctf1*$^{ti262}$ | GGTGAGTCAGTGTGGGGAAC | CCAGCAGGGCATGAAGTGAT |
| *b2m* | GCGGTTGGGATTTACATGTTG | GCCTTCACCCCAGAGAAAGG |
| *bad* | AGCAGCACCTCACTGTTCCT | CCAGTTTCCAGCAAGTCCTC |
| *bax* | GGAGATGAGCTGGATGGAAA | GGGCCACTCTGATGAAGACA |
| *bbc3* | GATGCCTTCAGCTTGGAC | GCCTGGACACTTCCTGTTCT |
| *bcl2* | GGATCGAGGAAAATGGAGGT | AAAACGGGTGGAACACAGAG |
| *bclxl* | CAACCATATTCAACCCTGGA | TTCTTGCGATTTCCTGCT |
| *bid* | ACCAGCGACCTACAGAGACC | TCTGCATTGACTGAAAGACCA |
| *bik* | TTGCTTCCACAGCTTCAAAA | ATGTAGTGCTGCGAGACCAG |
| *bim* | GCACTTTGATTTCCCTCAGC | TGGAGAAAGTCCGGTTCATC |
| *cdkn1a* | CAAGCCAAGAAGCGTCTAGTG | AACGGTGTCGTCTCTGGTTC |
| *cdkn2a/b* | CGAGGATGAACTGACCACAGC | CAACAGCCAAAGGTGCGTTAC |
| *hrpt1* | GAGGAGCGTTGGATACAGA | CTCGTTGTAGTCAAGTGCAT |
| *mdm2* | TGACAAAGAAACTGGTAAGA | AAACATAACCTCCTTCATGGT |
| *pmaip1* | ATGGCGAAGAAAGAGCAAAC | TCATCGCTTCCCCTCCATTTG |
| *tbp* | CAGGCAACACACCACTTTAT | AAGTTTACGGTGGACACAAT |
| *tp53* | TCCACTCTCCCACCAACATC | GGGAACCTGAGCCTAAATCC |
| *tp63* | CGGCCTGTTTGGACTATTTC | ACTCCATGATGCCTTTCCAG |

| | | |
|---|---|---|
| *tp73* | GGCCAATCCTCATCATCATC | TCCCTGAATGGTCTTCGTC |
| *Δ113tp53* | ATATCCTGGCGAACATTTGG | ACGTCCACCACCATTTGAAC |

**Supplementary Table 3. Percentage of HCC samples in the LIHC TCGA PanCancer Atlas with altered mRNA expression, Z-scores of genes encoding Nup107-160 complex components and DepMap portal gene essentiality CRISPR screens in human cancer cell lines.**

| Genes encoding Nup107-160 complex component | Percentage of samples with mRNA expression Z-score <-2 | Percentage of samples with mRNA expression Z-score >2 | Percentage of dependent cell lines in CRISPR screens |
|---|---|---|---|
| ***AHCTF1*** | 0.82 | 32.24 | 99.63 |
| *NUP37* | 0.00 | 6.01 | 14.36 |
| *NUP43* | 0.55 | 6.83 | 99.88 |
| *NUP85* | 0.00 | 18.31 | 100.00 |
| *NUP98* | 2.46 | 3.28 | 97.65 |
| *NUP107* | 0.27 | 7.92 | 96.29 |
| *NUP133* | 1.09 | 30.60 | 100.00 |
| *NUP160* | 0.00 | 5.46 | 97.77 |
| *SEC13* | 0.55 | 8.47 | 100.00 |
| *SEH1L* | 2.73 | 6.83 | 99.13 |

Heat-map based on percentage of hepatocellular carcinoma (LIHC) samples in the TCGA with mRNA expression levels of Nup107-160 components that are under-expressed (Z-score <-2: over 2 standard deviations below mean expression) or overexpressed (Z-score >2: over 2 standard deviations above mean expression) analysed using the cBioPortal for Cancer Genomics[39,40]. Genome-wide CRISPR screens, curated in the DepMap portal[48,49] reveals that the majority of Nup107-160 complex members are essential for the viability of human cancer cell lines.
